# Platelets promote acute liver injury via extracellular vesicles-mediated Aldolase A

**DOI:** 10.64898/2026.03.05.709856

**Authors:** Ruoxue Yang, Jinghua Liu, Kai Fu, Ting Wan, Yahui Li, Can Shen, Ling Yang, Keqin Wang, Zhao Shan

## Abstract

Platelets have emerged as active regulators of acute liver injury (ALI), yet the molecular mechanisms underlying their pathogenic functions remain poorly defined. Here, we demonstrate that platelets contribute to liver injury, in part, by metabolically reprogramming Kupffer cells (KCs). We show that platelets are actively recruited to the injured liver and communicate with KCs through extracellular vesicles (EVs), which deliver the glycolytic enzyme aldolase A (ALDOA) into recipient cells. This EV-mediated transfer promotes glycolytic reprogramming in KCs, contributing to their pro-inflammatory activation and amplification of hepatic injury. Genetic ablation of Aldoa in platelets attenuated liver injury, demonstrating a pathogenic role for platelet-derived ALDOA *in vivo*. Pharmacological inhibition of ALDOA with Aldometanib markedly attenuated liver injury. Circulating ALDOA levels were elevated in patients with ALI and correlated with disease severity. These findings uncover a platelet–macrophage metabolic axis that drives ALI and nominate ALDOA as a therapeutic target and biomarker.

## Introduction

Acute liver injury (ALI) is a life-threatening clinical syndrome characterized by rapid hepatocyte death, sterile inflammation, and loss of hepatic function, yet its cellular drivers remain incompletely defined (*Bernal et al., 2010; Lee, 2017*). ALI arises from diverse etiologies, including infections, toxins, and autoimmune disorders, with acetaminophen (APAP) overdose representing one of the most common causes (*Bernal et al., 2010; Lee, 2017*). A consistent clinical feature of patients with severe ALI is thrombocytopenia, manifested as a marked reduction in circulating platelet counts (*Hou et al., 2023; Tyagi et al., 2022*). In our previous study, we observed that these “lost” platelets accumulate within the liver, suggesting a previously unappreciated and potentially pathogenic role for platelets in ALI (*Shan et al., 2021*).

Platelets are anucleate blood components classically known for their role in hemostasis, but they are increasingly recognized as active modulators of inflammatory and immune responses (*Chauhan et al., 2020; Groeneveld et al., 2020; Hoeft et al., 2023; Levoux et al., 2021; Malehmir et al., 2019; Wong et al., 2013*). Nevertheless, technical challenges in platelet isolation, culture, and *in vivo* lineage tracing have limited mechanistic studies of platelet function in many diseases, including acute liver injury. Although platelet involvement in liver pathology has been suggested *(Miyakawa et al., 2015; Shan et al., 2021*), the precise molecular mechanisms by which platelets exert their effects remain poorly understood. Defining these pathways is essential for the development of targeted platelet-based therapeutic strategies.

Kupffer cells (KCs), the liver-resident macrophages strategically positioned within the hepatic sinusoids, serve as central orchestrators of hepatic immune responses *(Guilliams et al., 2022; Ponti et al., 2025*). Following liver injury, KCs integrate damage-associated signals and rapidly adopt dynamic functional states that can either exacerbate hepatocellular damage or promote tissue repair *(Deng et al., 2024; Triantafyllou et al., 2021*). How KC fate decisions are regulated within the injured hepatic microenvironment remains a key unresolved question. A defining feature of inflammatory macrophage activation is metabolic reprogramming, in which a shift toward aerobic glycolysis, reminiscent of the “Warburg effect,” fuels biosynthetic and pro-inflammatory programs in classically activated macrophages *(O’Neill & Pearce, 2016; Van den Bossche et al., 2017*). Whether a similar metabolic rewiring occurs in KCs during ALI, and how such changes might be imposed by non-immune cells such as platelets, has not been systematically explored.

Building on our previous discovery that platelets infiltrate the liver and physically interact with KCs (*Shan et al., 2021*), we hypothesized that platelet--KC crosstalk represents one mechanism by which KC function is reshaped, potentially exacerbating ALI. In this study, we sought to define the pathological consequences of platelet–KC interactions and to identify the critical molecular mediators involved. We demonstrate that platelets transfer the glycolytic enzyme aldolase A (ALDOA) to KCs via extracellular vesicles (EVs), thereby promoting glycolytic reprogramming in KCs and contributing to liver injury. Using platelet-specific Aldoa knockout mice, we establish the essential *in vivo* role of platelet-derived ALDOA. Moreover, pharmacological inhibition of ALDOA confers significant protection from liver injury, and circulating ALDOA levels are elevated in patients with ALI, underscoring the clinical relevance of our findings. Together, these results identify a platelet–KC metabolic axis that contributes to ALI pathogenesis and nominate ALDOA as a promising diagnostic biomarker and therapeutic target.

## Results

### Recruited platelets enhance glycolytic program in Kupffer cells

To investigate the role of platelets in APAP-induced liver injury (AILI), we established an AILI mouse model using wild-type C57BL/6J mice. Mice treated with APAP for 12 hours exhibited a significant increase in serum alanine aminotransferase (ALT) levels and extensive hepatic necrosis compared to PBS-treated controls, confirming the successful induction of the AILI model (*Figure 1-figure supplement 1A, B*). Immunofluorescence staining revealed a substantial infiltration of platelets into the liver following APAP injury (*Figure 1-figure supplement 1C*). Additionally, co-immunofluorescence staining of KCs and platelets revealed colocalization between these cell types, suggesting their interaction within the liver during AILI (*Figure 1-figure supplement 1D*). These findings are consistent with our previous study *(Shan et al., 2021*) and demonstrated that platelets are recruited to the liver and interact with KCs during AILI.

We next sought to characterize the response of KCs to APAP injury and determine whether these changes are influenced by platelets interactions. First, we isolated KCs from mice treated with APAP for 0 and 12 hours and performed RNA sequencing (RNA-seq). Kyoto Encyclopedia of Genes and Genomes (KEGG) pathway enrichment analysis showed a notable upregulation in glycolysis/gluconeogenesis pathways in KCs from mice treated with APAP for 12 hours compared to 0 hours, indicating a metabolic shift in KCs upon APAP treatment (Figure 1A). Consistently, heatmap analysis showed an overall increase in the expression of glycolysis-related enzymes in KCs at 12 hours (Figure 1B). The upregulation of glycolysis-related genes, including *Slc2a1*, *Pfkfb3*, and *Pkm*, at 12 hours post-APAP treatment was further validated by qRT-PCR (*Figure 1-figure supplement 1E*). In parallel, the expression of Hif1a and Il1β, markers associated with glycolytic and pro-inflammatory macrophage activation *(Chen et al., 2023*), was also significantly increased in KCs following APAP treatment (*Figure 1-figure supplement 1E*). Together, these findings suggest that KCs undergo a glycolytic and pro-inflammatory shift during early AILI.

**Figure 1.**
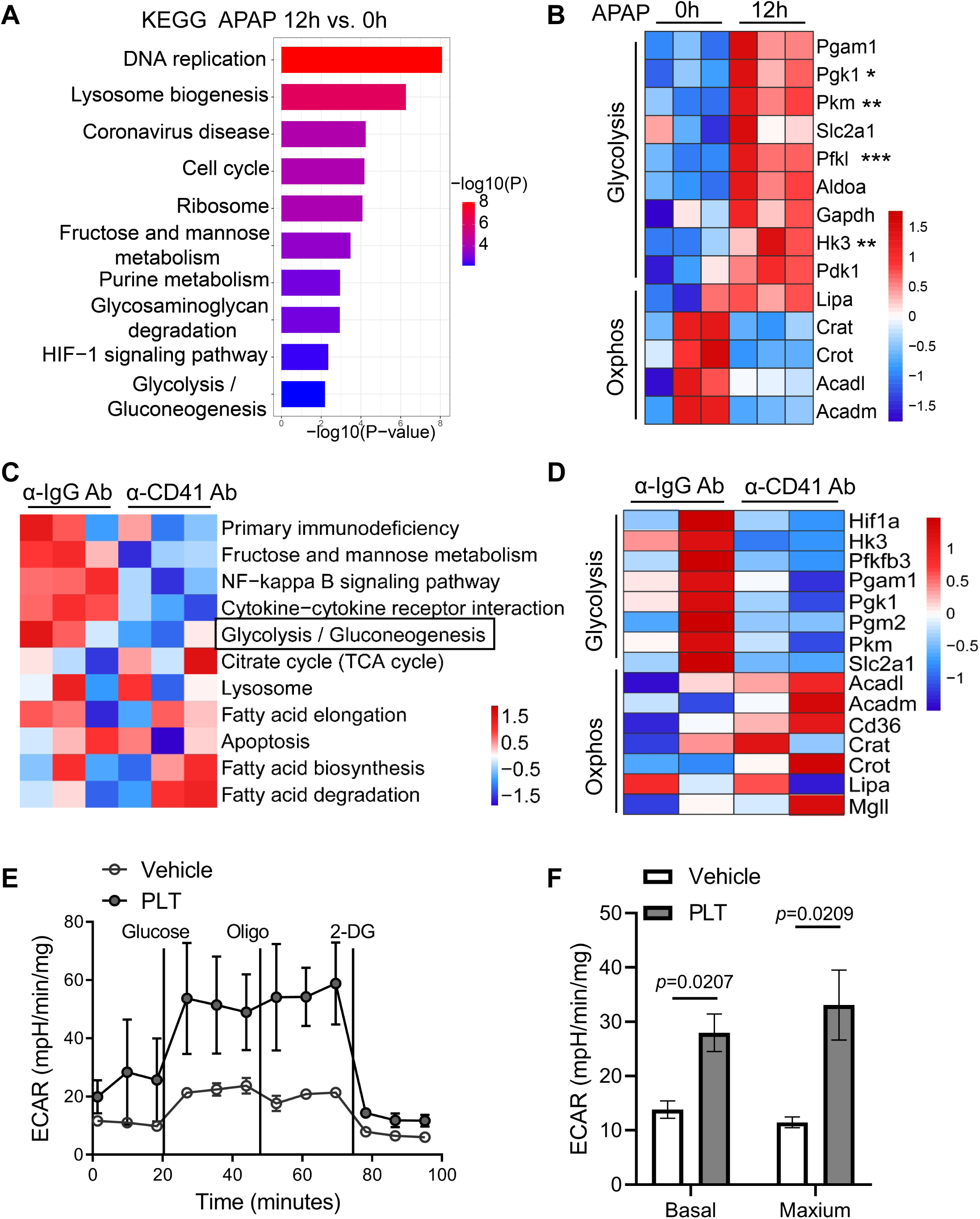
Recruited platelets enhance glycolytic program in Kupffer cells. **(A)** KEGG pathway enrichment analysis of differentially upregulated genes in Kupffer cells (KCs) isolated from mice treated with APAP for 12 hours versus 0 hours (n=3 mice/group), based on RNA-seq. **(B)** Heatmap depicting the expression of glycolysis and oxidative phosphorylation (Oxphos) genes in KCs following APAP treatment (n=3). *p < 0.05; **p < 0.01; ***p < 0.001 **(C)** Gene Set Variation Analysis (GSVA) of metabolic and immune pathways in KCs from mice treated with control IgG (α-IgG) or platelet-depleting (α-CD41) antibodies. The glycolysis pathway was significantly downregulated following platelet depletion (p=0.038). **(D)** Heatmap of glycolysis- and Oxphos-related gene expression in KCs from mice treated with α-IgG or α-CD41 antibodies followed by APAP for 12 hours. **(E)** Extracellular acidification rate (ECAR) of KCs cultured alone or co-cultured with mouse platelets for 2 hours, measured in response to glucose, oligomycin, and 2-deoxy-D-glucose (2-DG) (n=3). **(F)** Basal and maximal respiration of KCs described in (E). p-value was calculated using an unpaired two-tailed Student’ s t-test. Data are presented as mean ± SEM. *p* values are indicated; ns, not significant.

To determine whether recruited platelets were responsible for this metabolic reprogramming, we depleted platelets using an anti-CD41 antibody (α-CD41) prior to APAP challenge *(Figure 1-figure supplement 2A).* We then assessed the effects of platelet depletion on liver injury at 6 and 12 h after APAP administration. Immunofluorescence staining for CD41 confirmed efficient platelet depletion in the liver (*Figure 1-figure supplement 2B*). Notably, platelet depletion significantly attenuated liver injury at both time points, as evidenced by reduced necrotic areas and lower serum ALT levels (*Figure 1-figure supplement 2C--E*). Consistent with these findings, previous studies have shown that platelet depletion using anti-CD41 antibody also significantly reduced liver injury at 24 h after APAP challenge (*Miyakawa et al., 2015; Shan et al., 2021*). Collectively, these data support a protective effect of platelet depletion during the early phase of APAP-induced liver injury.

To further explore the effects of platelet depletion on the KC glycolytic program, we isolated KCs from platelet-depleted and control mice for RNA-seq analysis. Gene Set Variation Analysis (GSVA) of metabolic pathways in KCs from APAP-injured mice revealed that platelet depletion significantly reduced the glycolysis pathway (*p*=0.038) (Figure 1C). Consistent with this finding, heatmap analysis showed that the APAP-induced upregulation of glycolysis-related genes was attenuated in KCs from platelet-depleted mice, whereas the expression of oxidative phosphorylation (OXPHOS)-related genes was restored (Figure 1D). qRT-PCR further confirmed that platelet depletion significantly reduced the expression of glycolysis- and inflammation-related genes in KCs (*Figure 1-figure supplement 2F)*. Together, these findings demonstrate that platelets promote glycolytic reprogramming and a pro-inflammatory transcriptional state in KCs during acute liver injury.

To exclude the possibility that the protective effect of platelet depletion was attributable to altered APAP metabolism, we assessed hepatic GSH levels at 2 and 12 h after APAP administration. Hepatic GSH depletion was comparable between platelet-depleted and control mice at 2 h, with GSH levels recovering by 12 h in both groups (*Figure 1-figure supplement 2G)*. Consistent with these findings, hepatic CYP2E1 protein levels were also comparable between the two groups (*Figure 1-figure supplement 2H*). These results indicate that platelet depletion does not alter APAP bioactivation or its initial metabolic handling, suggesting that the protective effect of platelet depletion is unlikely to result from changes in APAP metabolism.

Finally, to establish a direct causal link, we isolated mouse platelets and co-cultured them with primary KCs in vitro. We measured the extracellular acidification rate (ECAR) in KCs cultured with either conditional medium (Vehicle) or isolated mouse platelets (PLT). Consistent with our in vivo findings, platelet-treated KCs exhibited significantly increased glycolytic activity (Figure 1E). Additionally, basal and maximal respiration rate was significantly increased in platelets-treated KCs, indicating elevated glycolytic activity (Figure 1F). Together, these findings demonstrate that platelet–KC interactions directly enhance glycolytic metabolism and alter the metabolic state of KCs in vitro.

### Kupffer cell glycolysis promotes APAP-induced liver injury

Having established that platelet-induced glycolysis in KCs is a prominent feature of the injury response, we next sought to determine whether this metabolic reprogramming functionally contributes to liver injury. To this end, we inhibited glycolysis with 2-deoxy-D-glucose (2-DG) during APAP challenge and used clodronate liposomes (CLDN) to determine whether the effects of glycolytic inhibition were dependent on KCs (Figure 2A). Flow cytometric analysis of hepatic non-parenchymal cells confirmed that CLDN pretreatment effectively depleted the CD11b^low^ F4/80^hi^ KC population, whereas control liposomes had no appreciable effect (*Figure 2-figure supplement 1A*). In KC-intact mice challenged with a low dose of APAP (210 mg/kg), 2-DG treatment significantly reduced both hepatic necrotic area and serum ALT levels compared with PBS-treated APAP controls (Figure 2B and 2C). Notably, this protective effect of 2-DG was largely lost following KC depletion, indicating that the protective effect of glycolytic inhibition was mediated, at least in part, through KCs. Consistent with our previous findings that KCs contribute to APAP-induced liver injury (*Shan et al., 2021*), KC depletion itself markedly attenuated hepatic necrosis and ALT elevation (Figure 2B and 2C).

**Figure 2.**
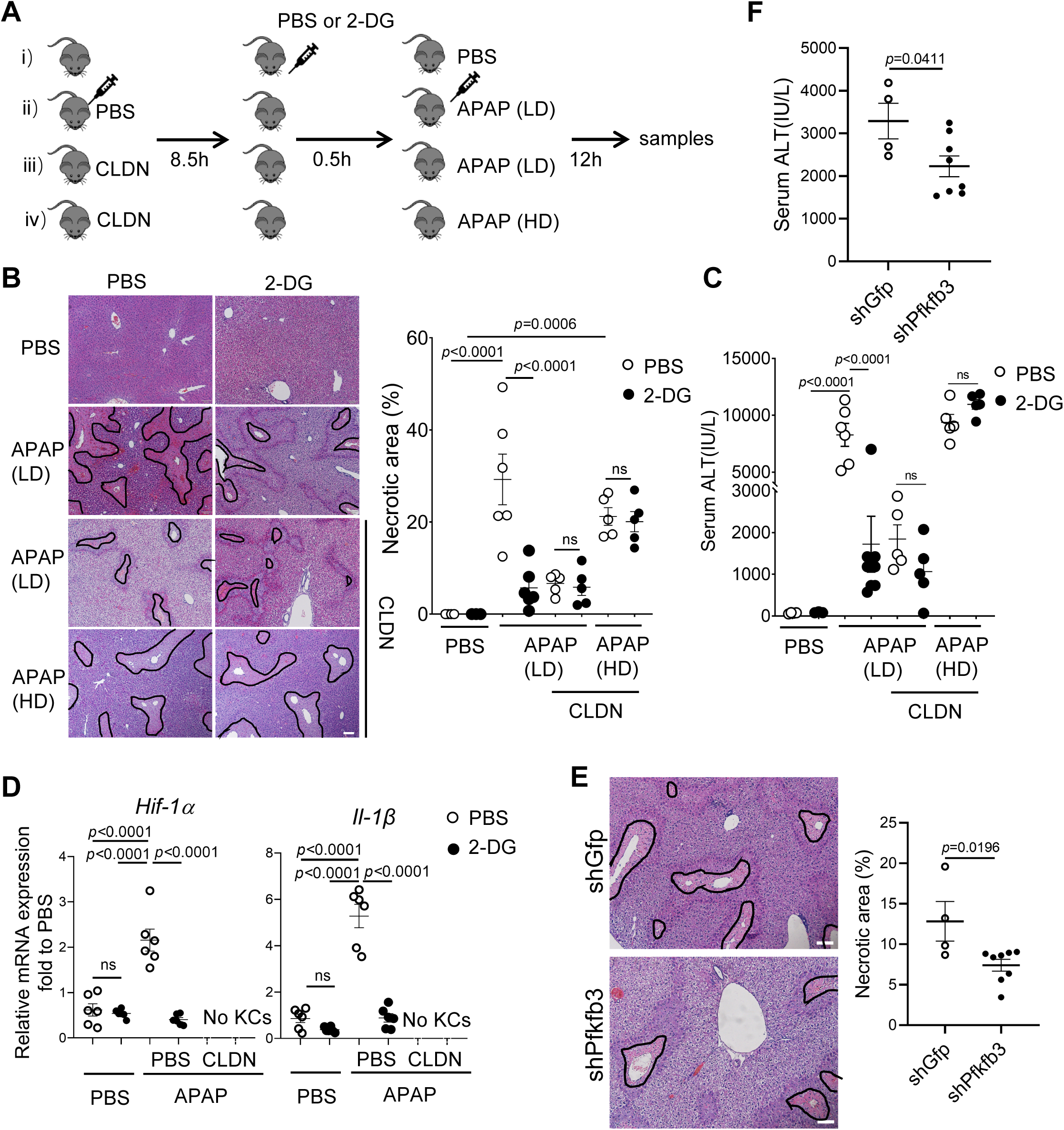
Kupffer cell glycolysis promotes APAP-induced liver injury. **(A)** Experimental timeline for assessing the role of Kupffer cell (KC) glycolysis in acetaminophen (APAP)-induced liver injury. Wild-type C57BL/6J mice were treated as follows: (i) Vehicle control: PBS or 2-deoxy-D-glucose (2-DG), followed 0.5 h later by PBS, with analysis 12 h later; (ii) KC-intact + APAP (210 mg/kg, low dose [LD]): control (PBS-loaded) liposomes for 8.5 h, followed by PBS or 2-DG for 0.5 h and then APAP challenge; (iii) KC-depleted + APAP (210 mg/kg, LD): clodronate liposomes (CLDN) for 8.5 h to deplete KCs, followed by PBS or 2-DG for 0.5 h and then APAP challenge; and (iv) KC-depleted + APAP (500 mg/kg, high dose [HD]): CLDN for 8.5 h, followed by PBS or 2-DG for 0.5 h and then APAP challenge. All mice were analyzed 12 h after APAP or PBS administration. n=6 mice/group. **(B)** Representative H&E-stained liver sections and quantification of the necrotic area in mice described in (A). Necrotic regions are outlined in black. Scale bars, 100 μm. **(C)** Serum alanine aminotransferase (ALT) levels in mice described in (A). **(D)** mRNA expression levels of inflammation-related genes in Kupffer cells (KCs) isolated from mice in (A), measured by qPCR. **(E)** Wild-type mice were intravenously injected with 6 × 10¹¹ genome copies (GC) of AAV-CD68-shPfkfb3 or control AAV-CD68-shGfp. Two weeks after AAV injection, mice were challenged with APAP (210 mg/kg, *i.p.*) for 12 hours. Representative H&E-stained liver sections from shGfp-and shPfkfb3-treated mice are shown. Necrotic regions are outlined in black. Scale bars: 100 µm. n=4 or 8 mice/group. **(F)** Serum alanine aminotransferase (ALT) levels in mice described in (E). Data are presented as mean ± SEM. Data in (B-D) were analyzed by one-way ANOVA. Data in (E, F) were analyzed using an unpaired Student’s *t*-test. *p* values are indicated; ns, not significant.

Because KC depletion markedly reduced liver injury following low-dose APAP, the remaining injury provided limited dynamic range for detecting an additional protective effect of 2-DG. We therefore challenged KC-depleted mice with a higher dose of APAP (500 mg/kg) to increase the injury burden and more rigorously assess whether glycolytic inhibition could confer additional protection in the absence of KCs. In KC-depleted mice challenged with high-dose APAP, extensive hepatic necrosis and robust ALT elevation were observed; however, 2-DG treatment did not further reduce either parameter (Figure 2B and 2C). Together, these findings indicate that the protective effect of 2-DG observed in KC-intact mice is largely dependent on the presence of KCs, supporting a functional contribution of KC glycolysis to APAP-induced liver injury. We next examined whether glycolytic inhibition altered the inflammatory phenotype of KCs. qPCR analysis of KCs isolated after APAP challenge showed that 2-DG significantly suppressed the expression of multiple inflammation-related genes that were strongly induced by APAP (Figure 2D). Thus, inhibition of glycolysis not only reduces tissue injury but also attenuates the inflammatory activation of KCs, further linking KC metabolic reprogramming to their injury-promoting inflammatory response.

To exclude the possibility that the protective effect of 2-DG and KC depletion were attributable to altered APAP metabolism, we assessed hepatic CYP2E1 expression and GSH levels. Hepatic CYP2E1 protein levels and NAPQI–protein adduct formation were comparable among all groups at 2 h after APAP administration, indicating that neither 2-DG treatment nor KC depletion altered the abundance of the major enzyme responsible for APAP bioactivation (*Figure 2-figure supplement 1B--C*). Consistent with this finding, baseline hepatic GSH levels were also comparable among groups, and APAP challenge resulted in a similar degree of GSH depletion at 2 h. By 12 h, hepatic GSH levels had largely recovered and were comparable among all groups (*Figure 2-figure supplement 1D*). Together, these results indicate that neither 2-DG treatment nor KC depletion significantly affects APAP bioactivation or the early kinetics of hepatic GSH depletion.

To further validate that KC-intrinsic glycolysis drives APAP injury, we pursued a complementary genetic loss-of-function approach. We delivered an adeno-associated virus serotype 8 (AAV8) encoding a short hairpin RNA (shRNA) against Pfkfb3, a rate-limiting glycolytic enzyme, under the control of the macrophage-specific CD68 promoter. Two weeks after transduction, KCs isolated from AAV-CD68-shPfkfb3-treated mice exhibited reduced Pfkfb3 mRNA expression compared to shGfp controls, confirming KC-restricted knockdown (*Figure 2-figure supplement 1E*). Although the knockdown efficiency in KCs was partial, this partial reduction was nonetheless sufficient to significantly attenuate APAP-induced injury, as indicated by markedly reduced hepatic necrosis (Figure 2E) and lower serum ALT levels (Figure 2F) in AAV-CD68-shPfkfb3-treated mice compared to shGfp controls following APAP challenge. Thus, genetic disruption of glycolysis specifically in macrophages phenocopied the protection afforded by pharmacological glycolytic inhibition.

### Platelets-derived extracellular vesicles promote glycolysis in Kupffer cells

We next sought to identify the mechanism by which platelets communicate with Kupffer cells to drive glycolysis. We collected supernatant from isolated mouse platelets after a 2-hour culture and co-cultured KCs with either conditional medium (Vehicle) or platelet supernatant (PLT sup) for 2 hours. KCs exposed to platelet supernatant exhibited increased expression of glycolysis-related and inflammatory response-related genes compared to the vehicle controls (Figure 3A), suggesting that platelets secrete soluble factors capable of inducing a metabolic switch in KCs. Previous studies have shown that platelets communicate with other cells through the release of EVs (*Mabrouk et al., 2023; Puhm et al., 2021*). To investigate whether EVs contribute to the effects of platelets on KCs, we isolated platelet-derived extracellular vesicles (PEVs) and, as a non-platelet EV control, hepatocyte-derived extracellular vesicles (HEVs) from AML12 cells. Western blot analysis detected the established EV marker ALG-2-interacting protein X (ALIX) in both PEVs and HEVs. CD63 was readily detected in PEVs but was not detectable in HEVs (Figure 3B). These findings confirmed the successful isolation of EV preparations from both platelet and hepatocyte sources and established HEVs as a non-platelet EV control for subsequent experiments.

**Figure 3.**
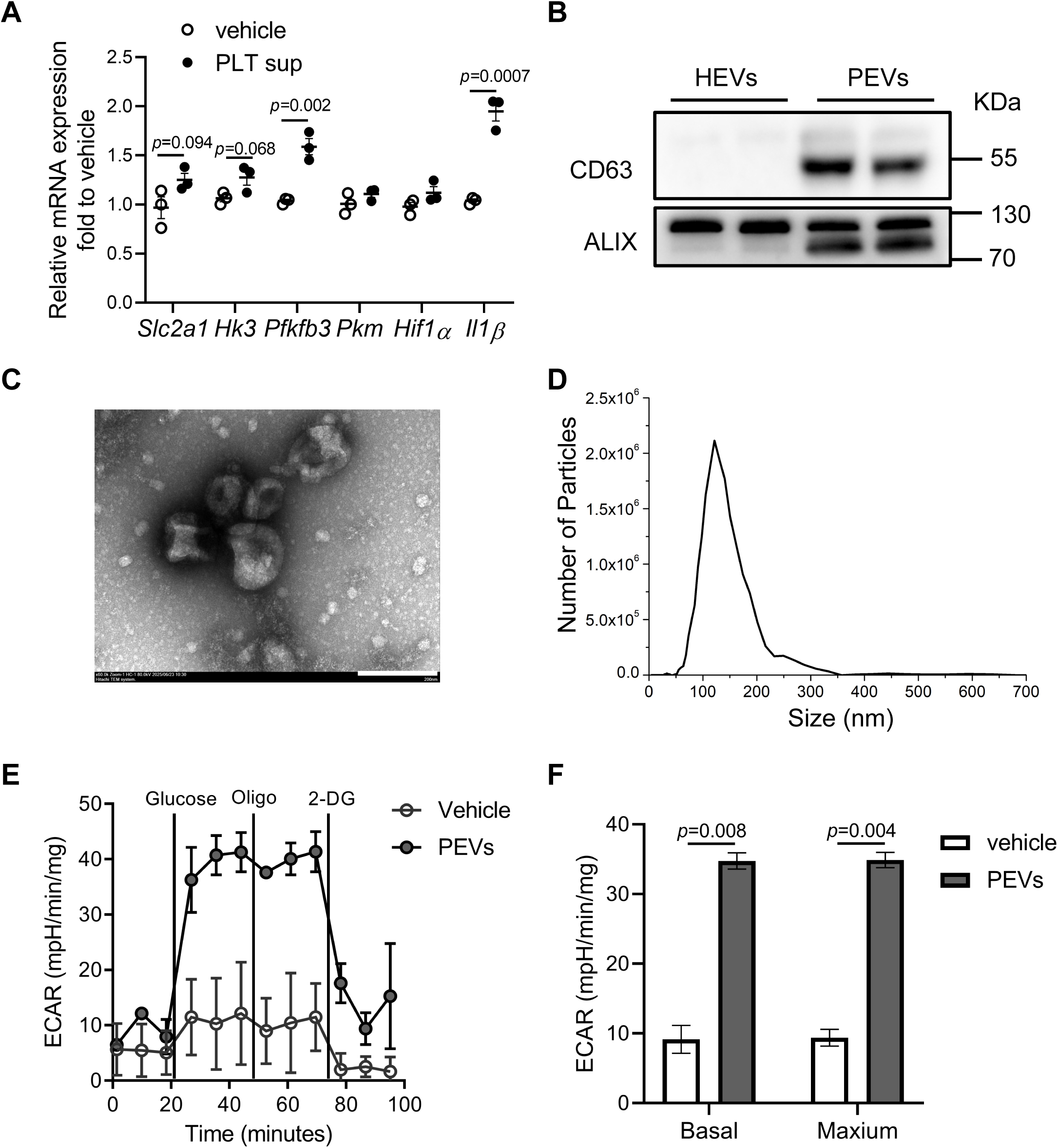
Platelets-derived extracellular vesicles promote glycolysis in Kupffer cells. **(A)** mRNA expression levels of glycolysis and inflammation-related genes in Kupffer cells (KCs) treated for 2 hours with supernatant from in vitro-cultured platelets (PLT sup), measured by qPCR. **(B)** Western blot analysis confirming the expression of extracellular vesicle (EV) markers (ALIX, CD63) in platelet-derived extracellular vesicles (PEVs). **(C)** Representative transmission electron microscopy (TEM) image of isolated PEVs. Scale bar, 200 nm. **(D)** Representative nanoparticle tracking analysis (NTA) profile of the isolated PEVs. **(E)** Extracellular acidification rate (ECAR) in KCs cultured alone or with PEVs for 2 hours, followed by sequential injection of glucose, oligomycin, and 2-deoxy-D-glucose (2-DG). **(F)** Basal and maximal respiration rates in KCs described in (E). Data are presented as mean ± SEM. Data in (A) and (F) were analyzed using an unpaired Student’s *t*-test. *p* values are indicated.

We next characterized the PEV preparations by transmission electron microscopy (TEM) and nanoparticle tracking analysis (NTA). TEM revealed spherical vesicles with a characteristic bilayer membrane structure (Figure 3C), while NTA showed that PEVs ranged predominantly from 59 to 288 nm in diameter, with a peak at approximately 105 nm (Figure 3D). To determine if these PEVs could directly modulate KC metabolism, we performed a Seahorse bioenergetic analysis. Kupffer cells co-cultured with isolated PEVs for just 2 hours exhibited a significantly elevated ECAR, indicating a robust increase in glycolytic flux (Figure 3E). Moreover, PEVs treatment also enhanced the basal and maximal respiratory capacity of the KCs (Figure 3F), confirming PEV’s effect on KC glycolysis. Together, these findings demonstrate that PEVs can directly promote glycolytic reprogramming in KCs, supporting platelet-derived EVs as one relevant source of EV-mediated metabolic regulation during AILI.

### Platelet-derived extracellular vesicles promote APAP-induced liver injury

Having established that PEVs can drive glycolysis in KCs in vitro, we next investigated their contribution to AILI in vivo. We used a rescue approach in platelet-depleted mice, which were treated with an anti-CD41 antibody to deplete platelets, followed by APAP administration together with vehicle, HEVs, or PEVs (Figure 4A). Immunofluorescence confirmed efficient depletion of platelets in the liver following anti-CD41 treatment (Figure 4B). As expected, platelet depletion protected mice from AILI, as evidenced by markedly reduced hepatic necrosis and serum ALT levels. Notably, administration of PEVs, but not HEVs, to platelet-depleted mice substantially restored APAP-induced liver injury at 12 h, as evidenced by significant increases in hepatic necrotic area and serum ALT levels (Figure 4C and 4D). We next examined whether PEVs and HEVs differentially affected KC metabolism in vivo. PEVs induced the expression of glycolysis-related genes in KCs, whereas HEVs did not produce a comparable increase in glycolytic gene expression, with the exception of Il1β (Figure 4E). Finally, to determine whether EVs are taken up by KCs in vivo, we intravenously injected PKH67-labeled PEVs or HEVs and analyzed their localization. Immunofluorescence staining revealed that a substantial number of TIM4-positive KCs had taken up both fluorescently labeled PEVs and HEVs within 12 h of administration (Figure 4F). Together, these findings suggest that PEVs are efficiently taken up by KCs and can promote their glycolytic reprogramming during AILI.

**Figure 4.**
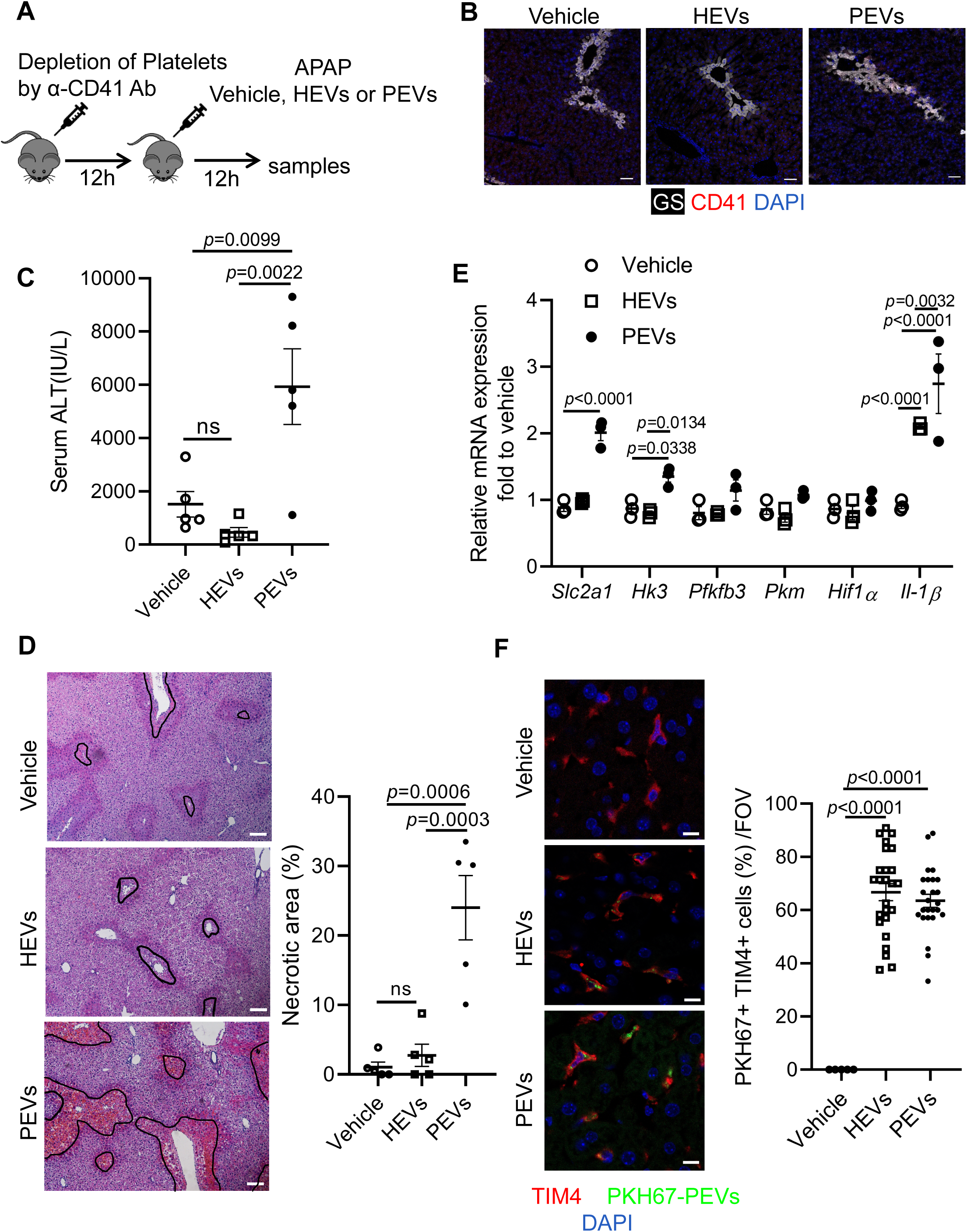
Platelet-derived extracellular vesicles promote APAP-induced liver injury. **(A)** Experimental timeline to assess the role of PEVs in APAP-induced liver injury following platelet depletion. Wild-type C57BL/6J mice were depleted of platelets using an anti-CD41 antibody (α-CD41 Ab) and then treated with Vehicle, HEVs or PEVs concurrently with APAP for 12 hours (n=5 mice/group). HEVs: extracellular vesicles derived from AML12 cells. **(B)** Immunofluorescence analysis confirming platelet depletion efficiency in liver sections from mice in (A), stained for CD41 (red), central vein (GS, white) and counterstained with DAPI (blue) for nuclei. Scale bar, 50 µm. **(C)** Serum alanine aminotransferase (ALT) levels from mice in (A). **(D)** Representative H&E-stained liver sections and quantification of necrotic area from mice in (A). Necrotic regions are outlined in black, and the percentage of necrotic area was quantified. Scale bars, 100 µm. **(E)** mRNA expression levels of glycolysis-related gene expression in KCs treated with vehicle, HEVs, or PEVs for 12 h, measured by qPCR. **(F)** Analysis of HEVs or PEVs uptake by Kupffer cells in vivo. Mice were intravenously (*i.v.*) injected with PKH67-labeled HEVs or PEVs for 12 hours. Liver sections were immunostained for Kupffer cells (TIM4, red) and nuclei (DAPI, blue). The number of PEVs-positive Kupffer cells was quantified. Scale bars, 10 µm. Data are presented as mean ± SEM. Data in (C-F) were analyzed by one-way ANOVA. *P* values are indicated.

### ALDOA from PEVs induces a glycolytic switch in Kupffer cells

To identify the specific component within PEVs responsible for driving glycolysis in KCs, we isolated PEVs and performed a quantitative mass spectrometry analysis of their protein cargo, which identified 121 proteins (Figure 5A, Supplementary File 1). To refine this list and select for proteins enriched in PEVs, we performed a comparative analysis with EVs derived from AML12 hepatocytes (PRIDE/ProteomeXchange: PXD084018). We excluded 53 proteins that were common to both PEVs and hepatocyte-derived EVs (HEVs), as well as cytoskeletal and coagulation-related proteins. This stringent filtering yielded a shortlist of seven candidate proteins: Aldolase A (ALDOA), Bridging integrator 2 (BIN2), Carboxylesterase 1C (CES1C), Inter-alpha-trypsin inhibitor heavy chain 2 (ITIH2), Murinoglobulin 1 (MUG1), Pleckstrin (PLEK), and Thrombospondin 1 (THBS1) (Figure 5A). Based on its central role as a catalytic enzyme in the glycolysis pathway, we selected ALDOA for further investigation. Western blot analysis revealed substantially higher levels of ALDOA in isolated PEVs than in HEVs, although a low level of ALDOA was also detectable in HEVs (Figure 5B). To test its functional capacity, we treated KCs with recombinant ALDOA (rALDOA) protein. Seahorse metabolic analysis revealed that rALDOA potently enhanced the glycolytic capacity of KCs, as shown by a marked increase in the ECAR (Figure 5C), as well as basal and maximal respiration rate (Figure 5D).

**Figure 5.**
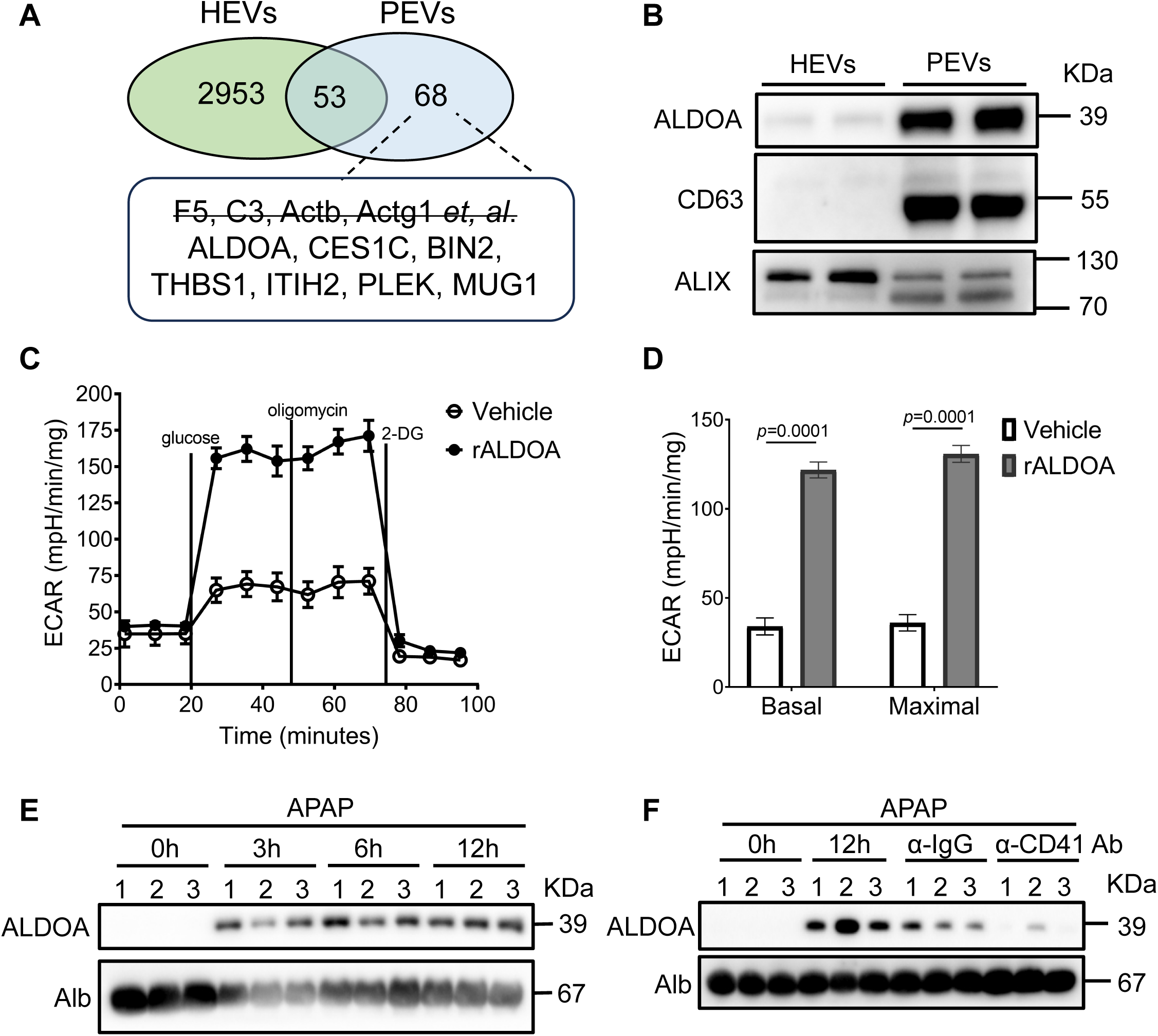
ALDOA from platelet-derived extracellular vesicles induces a glycolytic switch in Kupffer cells. **(A)** Schematic workflow for identifying PEV-derived protein mediators. Proteomic profiles of platelet-derived EVs (PEVs) and hepatocyte-derived EVs (HEVs) were compared. Proteins shared between PEVs and HEVs and those associated with the cytoskeleton or coagulation were excluded. Candidate PEV-derived protein mediators were defined as proteins detected in all three independent replicates with more than two unique peptides. **(B)** Western blot analysis of ALDOA expression in isolated PEVs and HEVs. CD63 and ALIX were used as EV markers. **(C)** Extracellular acidification rate (ECAR) in Kupffer cells (KCs) treated with vehicle or 500ng/ml recombinant ALDOA (rALDOA) for 2 hours, followed by sequential injection of glucose, oligomycin, and 2-deoxy-D-glucose (2-DG). Note that the rALDOA preparation contained a low concentration of imidazole from the protein purification buffer, which increased baseline glycolytic activity in KCs and may account for the higher basal ECAR observed in this experiment. **(D)** Basal and maximal respiration rates in KCs described in (C). **(E)** Western blot analysis of ALDOA expression in mouse serum at different time points (0, 3, 6, 12 h) after APAP treatment (n=3). **(F)** Western blot analysis of ALDOA expression in serum from mice under various conditions: untreated (0 h), 12 h post-APAP, and 12 h post-APAP following platelet depletion with either a control IgG antibody (α-IgG Ab) or an anti-CD41 antibody (α-CD41 Ab) (n=3). Data are presented as mean ± SEM. Data in (D) were analyzed by an unpaired two-tailed Student’s *t*-test. *p* values are indicated.

To establish the in vivo relevance of ALDOA during AILI, we measured its serum levels over time in AILI mouse models. Western blot analysis revealed a dynamic increase in circulating ALDOA, which was detectable as early as 3 hours post-APAP treatment and remained elevated at 6 and 12 hours (Figure 5E). To investigate whether platelets contribute to the increase in circulating ALDOA following APAP challenge, we depleted platelets prior to APAP administration. Platelet depletion resulted in a significant reduction in serum ALDOA levels compared with the control IgG-treated group (Figure 5F). Although this reduction may in part reflect the attenuated liver injury following platelet depletion, the finding is consistent with platelets contributing to the increase in circulating ALDOA during APAP-induced liver injury. Together, these findings support that platelet-derived vesicles represent an important source of ALDOA during liver injury. Upon delivery to KCs, platelet-derived ALDOA promotes a profound glycolytic switch in KCs.

### Platelet extracellular vesicles-mediated delivery of Aldoa promotes APAP-induced liver injury

To establish the pathophysiological role of platelet-derived ALDOA in vivo, we generated platelet-specific Aldoa knockout mice (hereafter, Aldoa-PtKO mice). Efficient deletion of Aldoa was confirmed by genotyping and Western blot analysis of platelet lysates (*Figure 6-figure supplement 1A--B*). Following APAP challenge, Aldoa-PtKO mice exhibited significantly less liver injury than Aldoa^fl/fl^ littermate controls, as evidenced by reduced serum ALT levels and a smaller hepatic necrotic area (Figure 6A and 6B). We next asked whether the protection conferred by platelet ALDOA deficiency resulted from altered APAP metabolism. Hepatic CYP2E1 protein levels and NAPQI–protein adduct formation were comparable between Aldoa-PtKO and control mice (*Figure6-figure supplement 2A--B*), indicating that platelet ALDOA deficiency does not affect APAP bioactivation. Consistent with these findings, hepatic GSH levels showed a similar and profound depletion at 2 h after APAP challenge in both genotypes, followed by complete recovery at 6 and 24 h (*Figure 6-figure supplement 2C*). Thus, the protective effect of platelet ALDOA deficiency is unlikely to result from altered APAP bioactivation or GSH depletion. We next examined downstream consequences of APAP-induced injury. Dihydroethidium (DHE) staining of liver sections at 24 h after APAP challenge revealed significantly reduced ROS accumulation in Aldoa-PtKO mice compared with control mice *(Figure 6-figure supplement 2D*). In parallel, KCs isolated from APAP-challenged Aldoa-PtKO mice showed reduced expression of glycolysis- and inflammation-related genes (*Figure 6- figure supplement 2E*). Together, these findings indicate that platelet-derived ALDOA promotes APAP-induced liver injury downstream of APAP bioactivation, in association with enhanced glycolytic and inflammatory reprogramming of KCs and increased hepatic oxidative stress.

**Figure 6.**
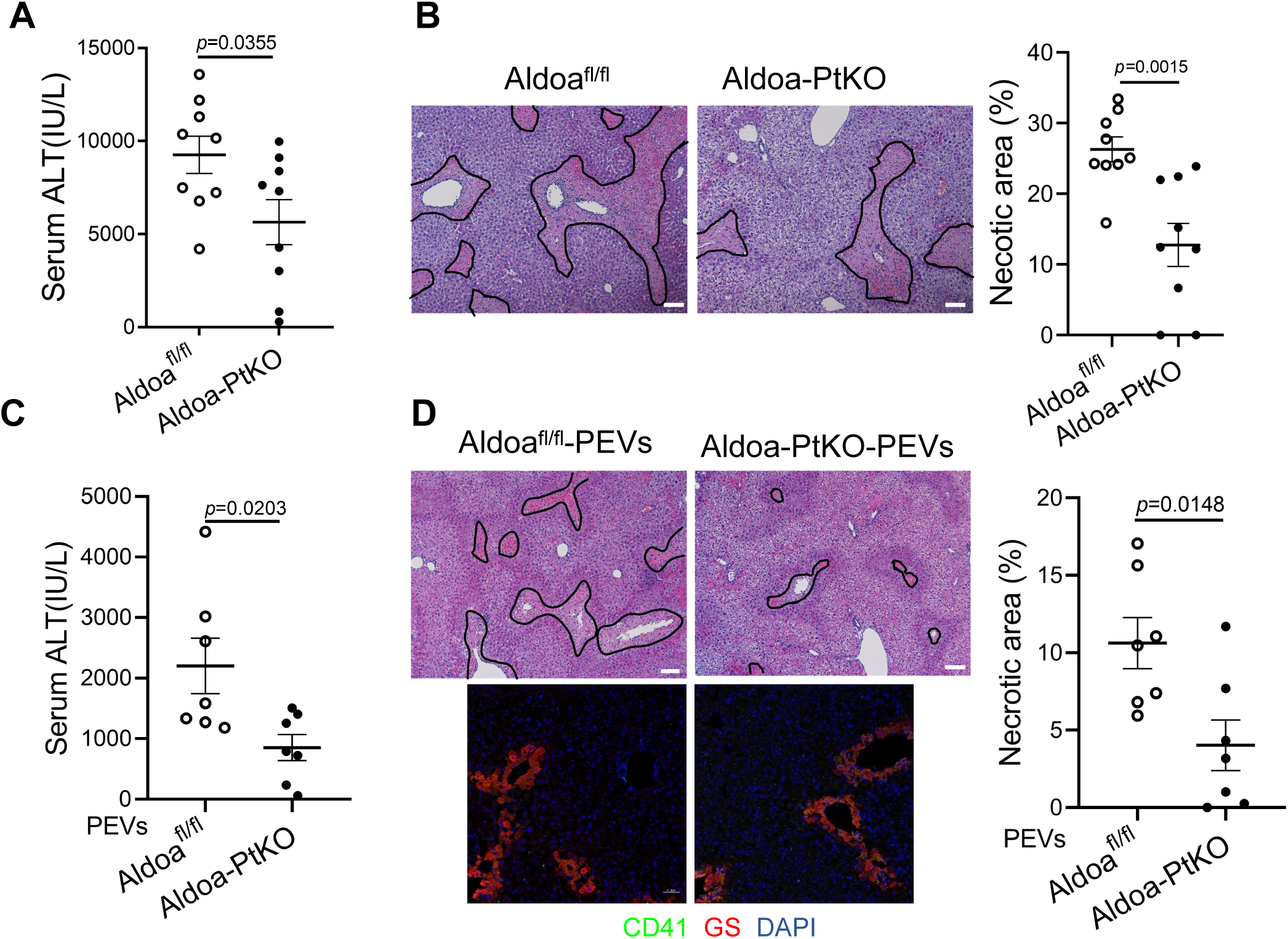
Platelet extracellular vesicles-mediated delivery of Aldoa promotes APAP-induced liver injury. **(A)** Serum ALT levels were measured in *Aldoa^fl/fl^* and Aldoa-PtKO mice treated with 210mg/kg APAP for 24 hours. n=9 mice/group. **(B)** Hematoxylin & Eosin (H&E) was performed to evaluate liver morphology in liver sections of mice in (A) (left). Necrotic areas were outlined in black, and the percentage of necrotic area was quantified (right). **(C)** Wild-type C57BL/6J mice were depleted of platelets using an anti-CD41 antibody (α-CD41 Ab) and then treated with PEVs from either *Aldoa^fl/fl^* or Aldoa-PtKO mice concurrently with APAP for 12 hours. Serum ALT levels were measured. n=7 mice/group. **(D)** Hematoxylin & Eosin (H&E) was performed to evaluate liver morphology in liver sections of mice in (C) (upper panel). Scale bar, 100 µm. Necrotic areas were outlined in black, and the percentage of necrotic area was quantified (right panel). Immunofluorescence analysis confirming platelet depletion efficiency in liver sections from mice in (C) (lower panel), stained for CD41 (red), central vein (GS, white) and counterstained with DAPI (blue) for nuclei. Scale bar, 50 µm. Data are presented as mean ± SEM. *p*-values were calculated using an unpaired two-tailed Student’s *t*-test (A, B, D). *p* values are indicated.

To further establish whether ALDOA acts as a pathogenic cargo of PEVs, we generated PEVs from Aldoa^fl/fl^ and Aldoa-PtKO mice. Western blot analysis confirmed that ALDOA was undetectable in PEVs from Aldoa-PtKO mice, whereas the EV marker TSG101 was comparable between genotypes (*Figure 6-figure supplement 2F*), indicating comparable recovery of PEVs. We next depleted endogenous platelets in wild-type mice using an anti-CD41 antibody and reconstituted the mice with equal amounts of PEVs derived from either Aldoa^fl/fl^ or Aldoa-PtKO mice, followed by APAP challenge. Platelet depletion was confirmed by CD41 immunofluorescence staining (Figure 6D, bottom panel). Notably, mice receiving ALDOA-deficient PEVs exhibited significantly lower serum ALT levels and reduced hepatic necrosis compared with mice receiving control PEVs (Figure 6C and 6D, top panel). Together, these findings provide direct in vivo evidence that PEV-associated ALDOA contributes to APAP-induced liver injury.

### ALDOA serve as a potential therapeutic target for acute liver injury

We next investigated the therapeutic potential of pharmacologically inhibiting ALDOA after the initiation of injury. Administration of the ALDOA inhibitor Aldometanib *(Zhang et al., 2022*) one hour after APAP overdose (Figure 7A) significantly attenuated liver injury, as evidenced by reduced serum ALT levels and hepatic necrosis (Figure 7B and 7C). This protective effect was not attributable to altered APAP bioactivation, as hepatic CYP2E1 protein levels and NAPQI–protein adduct formation were comparable between Aldometanib-treated and control mice (*Figure 6-figure supplement 2G*). We next asked whether this pathway also contributes to other forms of acute liver injury. In a Concanavalin A (Con A)-induced immune-mediated liver injury model, KC depletion similarly attenuated hepatic injury (*Figure 7-figure supplement 1A--B*). Importantly, Aldometanib treatment also significantly reduced Con A-induced liver injury (*Figure 7- figure supplement 1C--E*), indicating that ALDOA-driven pathogenic signaling extends beyond APAP-induced liver injury to immune-mediated hepatitis.

**Figure 7.**
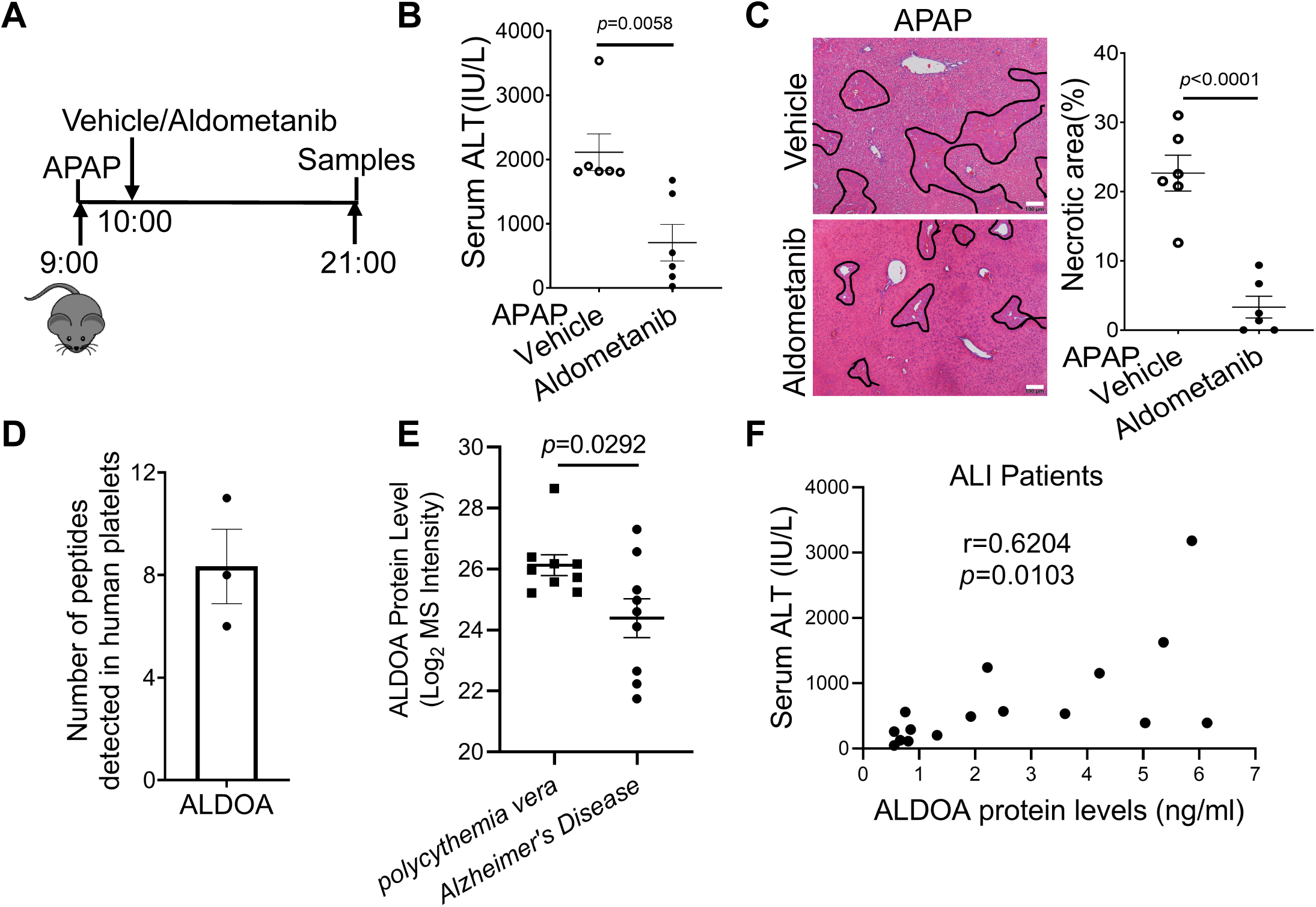
ALDOA serve as a potential therapeutic target for acute liver injury. **(A)** Experimental strategy for inhibiting ALDOA in vivo during AILI. Wild-type C57BL/6J mice were starved overnight (5 PM-9 AM), followed by an intraperitoneal injection of APAP at 9 AM. Mice were treated with 2 mg/kg Aldometanib one hour after APAP injection, and samples were collected 12 hours post-APAP. n=6 mice/group. **(B)** Serum ALT levels were measured in mice described in (A). **(C)** H&E staining was performed to evaluate liver morphology in liver sections of mice described in (A). Necrotic areas were outlined in black, and the percentage of necrotic area was quantified. **(D)** Platelets were isolated from healthy donors, and proteins were identified in the alpha granule fractions using liquid chromatography-tandem mass spectrometry (LC-MS/MS) as described by Maynard et al., 2007. **(E)** Extracellular vesicles (EVs) were isolated from serum of patients with polycythemia vera (*Fel et al*. (2019)) or Alzheimer’s Disease (Nielsen et al. (2021)). Protein content within the EVs was characterized using liquid chromatography–tandem mass spectrometry (LC–MS/MS). Among the identified proteins, ALDOA intensity was quantified. **(F)** The correlation between ALDOA protein levels and serum ALT levels in patients with ALI (the detailed patients information is in Supplementary File 2). Correlations were evaluated by the Pearson correlation coefficient. n=16. Data are presented as mean ± SEM. *p*-values were calculated using an unpaired two-tailed Student’s *t*-test (A, C). *p* values are indicated.

Finally, we sought to assess the clinical relevance of these findings in human disease. Reanalysis of a published proteomic dataset confirmed that ALDOA is present in human platelet α-granules (*Maynard et al., 2007*) (Figure 7D). Consistent with this finding, analysis of serum-derived EVs revealed significantly higher EV-associated ALDOA levels in patients with polycythemia vera (*Fel et al., 2019*), a disorder associated with hepatic involvement, than in patients with Alzheimer’s disease *(Nielsen et al., 2021*) (Figure 7E). Most importantly, in a cohort of patients with ALI, circulating ALDOA levels showed a strong positive correlation with serum ALT levels, a clinical indicator of hepatocellular injury (Figure 7F). Together, these findings identify ALDOA as a therapeutically actionable mediator of acute liver injury and suggest that circulating ALDOA may serve as a potential biomarker of disease severity.

## Discussion

The progression of AILI is a complex process involving not only the initial hepatocyte necrosis caused by NAPQI but also a robust sterile inflammatory response that amplifies the damage (*Ben-Moshe et al., 2022*). While the role of immune cells like KCs has been acknowledged, the specific signals that activate them remain incompletely understood. In this study, we elucidate a novel and critical axis in which recruited platelets, via the release of EVs loaded with the glycolytic enzyme ALDOA, drive a pathogenic metabolic and inflammatory reprogramming of KCs, thereby contributing to the exacerbation of liver injury. Furthermore, we identify ALDOA as a viable therapeutic target for ALI (Figure 8).

**Figure 8.**
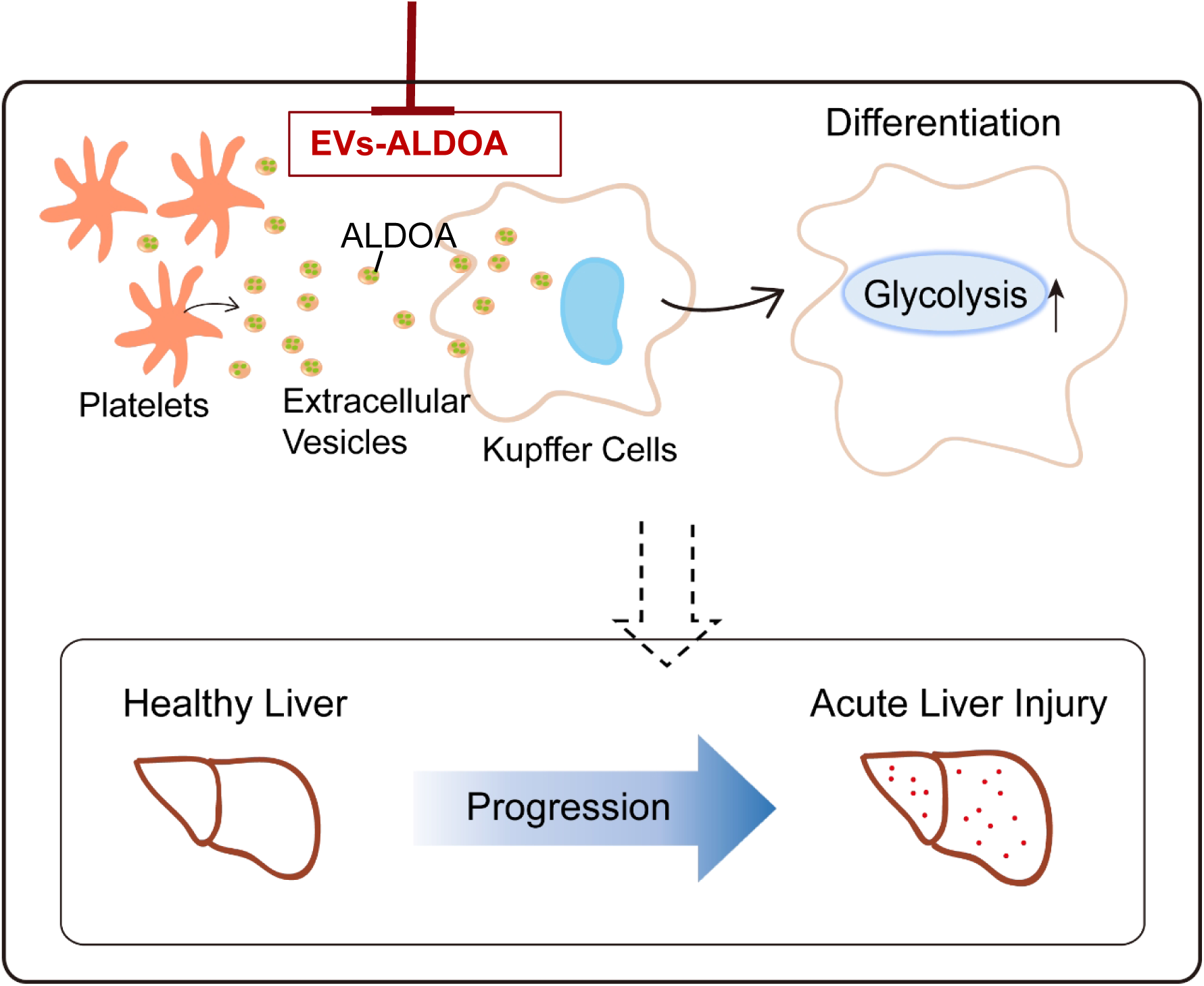
Platelets promote acute liver injury by transferring ALDOA to Kupffer cells via extracellular vesicles. Platelet-derived extracellular vesicles (PEVs) deliver Aldolase A (ALDOA) to Kupffer cells, enhancing their glycolytic activity and driving acute liver injury. Targeting the ALDOA pathway in platelets represents a promising therapeutic strategy.

Our findings extend previous observations that platelets are rapidly recruited to the liver following APAP overdose and physically interact with KCs (*Shan et al., 2021*). More importantly, complementary pharmacological inhibition with 2-DG and genetic suppression using AAV-mediated shRNA provided additional evidence that KC-intrinsic glycolysis contributes to injury amplification. Although AAV-mediated shRNA achieved only partial suppression in tissue-resident macrophages, this reduction suggests significant protection, further supporting a functional role for KC glycolysis in AILI. Although our study focused specifically on platelet–KC crosstalk and did not examine metabolic effects of platelets on other hepatic cell populations, our findings identify platelet–KC metabolic communication as one mechanism through which platelets can modulate the inflammatory response during sterile liver injury. This interpretation does not exclude additional, KC-independent effects of platelets on other hepatic cell populations.

The central mechanistic insight of our study is that PEVs serve as key mediators of platelet-KC communication. Our findings identify PEVs as an important mediator of platelet–KC communication. PEV administration was sufficient to restore liver injury in platelet-depleted mice, providing direct evidence for their pathogenic activity. Adoptive transfer of isolated PEVs substantially restored liver injury in platelet-depleted mice, providing direct evidence for their pathogenic role. This finding adds a new dimension to the understanding of platelet function in sterile inflammation, positioning them not only as coordinators of coagulation and inflammation but also as active regulators of cellular metabolism in recipient cells via vesicular communication *(Mabrouk et al., 2023; Puhm et al., 2021*).

Our proteomic and functional analyses identified ALDOA as a key protein cargo of PEVs that promotes glycolytic reprogramming in KCs. This finding is particularly intriguing because ALDOA is a canonical intracellular glycolytic enzyme, whereas its presence in the extracellular space and within EVs suggests a potential non-canonical role in intercellular communication. We found that extracellular ALDOA was sufficient to enhance glycolytic activity and the expression of inflammatory genes in KCs. The increase in circulating ALDOA during AILI further suggests that extracellular ALDOA may participate in the response to liver injury, although its cellular sources in vivo are likely to be heterogeneous. Given that PEVs are efficiently taken up by KCs in vivo, our findings raise the possibility that PEV-associated ALDOA is delivered into KCs through EV uptake, where it may contribute directly to glycolytic reprogramming. Thus, platelet-derived EVs may provide a mechanism for delivering an otherwise intracellular glycolytic enzyme to KCs and thereby linking platelet activation to metabolic and inflammatory responses during AILI.

The translational potential of targeting this pathway is strongly supported by our genetic and pharmacological studies. The protection observed in platelet-specific Aldoa knockout mice (Aldoa-PtKO) confirms the in vivo relevance of our mechanistic model. Crucially, the protective effect of the ALDOA inhibitor Aldometanib, even when administered after APAP overdose, highlights a promising therapeutic window for intervention that targets the secondary inflammatory wave rather than the initial metabolic insult. This feature may provide a complementary therapeutic opportunity by targeting the secondary inflammatory amplification of injury beyond the early window in which N-acetylcysteine is most effective (*Andrade et al., 2019*). The efficacy of Aldometanib in both APAP- and Con A-induced liver injury models suggests that ALDOA may represent a shared pathogenic pathway across distinct forms of acute liver injury.

Finally, the clinical relevance of our findings is underscored by the analysis of human samples. The presence of ALDOA in human platelet granules and its elevated levels in serum EVs from patients with liver-involving diseases like polycythemia vera provide a crucial bridge from mouse models to human pathology. Most importantly, the strong positive correlation between circulating ALDOA levels and serum ALT in patients with ALI highlights ALDOA as a potential biomarker of disease severity and further supports its translational potential as a therapeutic target.

In conclusion, this study identifies a previously unrecognized platelet–EV–ALDOA metabolic pathway: Platelets → PEVs → ALDOA → KC Glycolytic Switch → Inflammatory Amplification → Hepatocyte Necrosis. This pathway operates alongside other known mechanisms of liver injury. By identifying ALDOA as an important effector of this axis, our work opens up new avenues for therapeutic intervention. Targeting the ALDOA-mediated metabolic communication between platelets and KCs could represent a novel and effective strategy for treating ALI in patients.

## Materials and Methods

### Human sera

Serum samples from healthy volunteers and patients diagnosed with acute liver injury were collected at the Affiliated Hospital of Yunnan University (AHYNU), Kunming, China. The study was designed and carried out in accordance with the principles of AHYNU and approved by the Ethics Committee of AHYNU (Approval number 2024351). All patients provided written informed consent. Serum ALDOA levels in acute liver injury patients were quantified by western blot analysis, using 2 ng of recombinant ALDOA (rALDOA) as a standard, and band intensities were analyzed with ImageJ software.

### Animal experiments and procedures

C57BL/6J mice (Strain No. N000013) were purchased from GemPharmatech. Aldoa^fl/fl^ (Strain No. NM-KO-226330) were purchased from Southern Model Biotechnology Co., Ltd. To achieve platelet-specific deletion, Aldoa^fl/fl^ mice were crossed with PF4-cre transgenic mice (Jackson Laboratory, Stock #008535), in which Cre recombinase expression is driven by the platelet factor 4 promoter. PF4-Cre; Aldoa ^fl/fl^ (Aldoa-PtKO) mice and their Aldoa^fl/fl^ littermates were used as controls. Genotyping was performed by PCR on genomic DNA extracted from tail clips using the following primers: Aldoa ^fl/fl^: F-GTTAGGCAAGCGCTCTGTTG, R- GTCAGCTTTATGGCCCCTGT. PF4-Cre: F1-CCAAGTCCTACTGTTTCTCACTC, R1- TGCACAGTCAGCAGGTT, F2-CTAGGCCACAGAATTGAAAGATCT, R2- GTAGGTGGAAATTCTAGCATCATCC. All the primers are from 5’end to 3’ end.

All mouse colonies were maintained at the Laboratory Animal Center of Yunnan university. Animal studies described have been approved by the Yunnan University Institutional Animal Care and Use Committee (IACUC, Approval No. YNU20220262 for C57BL/6J strain, Approval No. YNU20240898 for Aldoa^fl/fl^ strain, Approval No. YNU20220387 for PF4-Cre strain). For APAP treatment, mice (8-12 weeks old) were fasted overnight (5:00pm to 9:00am) before *i.p.* injected with APAP (Sigma, Cat #A7085) at a dose of 210 mg/kg for male mice. Male mice have been the choice in the vast majority of the studies of AILI reported in the literature (Ben-Moshe et al., 2022; Matchett et al., 2024). Therefore, we used male mice in the majority of the experiments presented. However, we observed a similar phenotype in female mice as in male mice regarding the interaction between KCs and platelets(Shan et al., 2021). For Concanavalin A treatment, mice were *i.v.* injected with Saline or Concanavalin A (Sigma, Cat #C2010) at a dose of 15mg/kg. Liver paraffin sections and sera were harvested at time points indicated in the figure legends. H&E staining and ALT measurement (Teco Diagnostics, Cat #A524-150) were performed to examine liver injury.

Platelet depletion: WT mice were *i.v.* injected with anti-IgG (BD Pharmingen, Cat #553922, 2mg/kg) or anti-CD41 antibody (BD Pharmingen, Cat #553847, 2mg/kg) to deplete platelets at 12 hours before APAP treatment, followed by 210mg/kg APAP *i.p.* treatment for additional another 12 hours.

Kupffer cells depletion: WT mice were *i.v.* injected with 100 μL Clodronate liposomes (CLDN) to deplete Kupffer cells 9 hours before APAP or Con A treatment, followed by 210mg/kg APAP *i.p.* treatment or Con A treatment.

Inhibition of glycolysis: WT mice were *i.p*. injected with 250mg/kg 2-DG (Sigma, Cat# 8375, dissolved with PBS at a concentration of 50mg/mL) half an hour before APAP treatment, followed by 210mg/kg APAP *i.p.* treatment for additional another 12 hours.

Blocking endogenous ALDOA: WT mice were *i.p*. injected with 2mg/kg Aldometanib (MCE, Cat# HY-148189, dissolved with N-Methyl-2-pyrrolidone (Sangon Biotech, Cat# 872-50-4) at a stock concentration of 50mg/mL, diluted with PBS) one hour after APAP treatment, liver paraffin sections and sera were harvested at 12 hours after APAP treatment.

EVs labeling and injection: PEVs were isolated from mouse platelets culture supernatant by ultracentrifuge. HEVs were isolated from normal AML12 cells supernatant. And stained with PKH67 Green Fluorescent (sigma, Cat #MINI67) in 37°Cfor 30min. EVs were washed with PBS for 1 time. And then collects EVs by ultracentrifuge. WT mice were *i.v.* injected with anti-CD41 antibody (BD Pharmingen, Cat #553847, 2mg/kg) to deplete platelets at 12 hours before APAP treatment, followed by 210 mg/kg APAP *i.p.* treatment and 50 μg EVs (PEVs from wild-type mice, Aldoa^fl/fl^, Aldoa-PtKO mice or HEVs from ALM12 cells) (per mouse) *i.v.* treatment at the same time for additional another 12 hours.

Adeno-Associated Virus (AAV)-Mediated Gene Transfer: For knockdown Pfkfb3, the following Adeno-associated virus serotype 2/8 (AAV2/8) under a macrophage-special CD68 promoter was used in this study as previously reported(Zou et al., 2024). 6-week- old wild-type mice were intravenously injected with 6 × 10^¹¹^ genome copies (GC) of AAV-

CD68-shPfkfb3 or control AAV-CD68-shGfp. Two weeks later, mice were fasted overnight (5:00pm to 9:00am) and *i.p.* injected with 210 mg/kg APAP for 12 hours. For confirming KC-specific knockdown efficiency, naïve Kupffer cells were isolated, and Pfkfb3 mRNA expression was measured by qPCR.

### Preparation of Kupffer cells

Wild-type C57BL/6J mice were anesthetized using a CO_2_ chamber. To avoid contamination of the peritoneal cavity with fur, the skin and peritoneum were opened separately. The inferior vena cava was clamped with forceps, and a catheter was inserted into the superior vena cava via the right atrium. The portal vein was then cut once the buffer started to perfuse into the liver. The liver was perfused with 1× Hank’s Balanced Salt Solution (HBSS) containing 53.6 mM KCl, 4.4 mM KH_2_PO_4_, 1367.5 mM NaCl, 3.36 mM NaHPO_4_, 55.5 mM D-Glucose, and 100 mL 1 M HEPES (pH 7.2-7.4) for 3-5 minutes until it appeared clear. Subsequently, the buffer was switched to a collagenase digestion solution (1×HBSS with 1.25 mM MgSO_4_, 4 mM CaCl_2_, Collagenase Type I, and 50 μg/mL DNase) for 15-20 minutes of digestion. Post-digestion, the liver was dissected and placed in chilled 1×HBSS in a Petri dish to keep it cool until further processing. The gallbladder was removed, and the liver was chopped into 2-3 mm pieces and incubated in 10-20 mL digestion buffer at 37°C for 15 minutes. The liver tissue was then mashed using a 70 μm cell strainer placed over a 50 mL tube and rinsed with 1×HBSS to a final volume of 50 mL. The suspension was centrifuged at 50×g for 5 minutes (acceleration/brake = 0) to remove hepatocytes, with the resulting supernatant containing nonparenchymal cells (NPCs). The supernatant was further centrifuged at 450 × g for 5 minutes (acceleration/brake ≥ 5), and the pellet, containing hepatic NPCs, was resuspended in 4 mL of 20% OptiPrep solution diluted with 1×HBSS. This suspension was carefully layered with 5 mL of 1×HBSS buffer in a 15 mL tube and centrifuged at 3000 rpm for 17 minutes (acceleration/brake = 0). The cell fraction between the 20% OptiPrep and 1×HBSS layers, containing KCs and liver endothelial sinusoidal cells (LESCs), was collected and washed with 1×HBSS.To isolate KCs, the cell fraction was incubated in a cell culture dish at 37°C for 10 minutes. KCs adhered to the plate, while LESCs remained in suspension. The adhered KCs were then collected for subsequent experiments. The purity and viability of isolated KCs reached 90% and 80%, respectively, which was confirmed by staining with anti-Timd4 antibodies or trypan blue.

### Kupffer cells cultivated in vitro

KCs culture medium consisted of DMEM (VivaCell, Cat #C3113-0500) supplemented with 10% (vol/vol) fetal bovine serum (FBS, VivaCell, Cat #C04001-500), 100 IU penicillin, and 100 μg/mL streptomycin. KCs were cultured for 2 hours. For experiments involving KCs, 1×10^^6^ cells seeded in one well of a 12-well plate were co-cultured with either 0.5×10^^6^ isolated mouse platelets or vesicles isolated from 1×10^^6^ isolated mouse platelets for 2 hours. After co-culture, the cells underwent various experiments including Seahorse assay, and RT-PCR. In another study, KCs were treated with platelet supernatant for 2 hours. Platelet supernatant was collected from isolated mouse platelets after a 2-hour culture. In additional experiments, KCs were treated with recombinant ALDOA for 12 hours. In some experiments, cells were treated with either vehicle or 5 μg/mL PEVs for 2 hours.

### Isolation of mouse platelets

Blood was collected from CO_2_-sacrificed mice via cardiac puncture using a 1/10 volume of ACD buffer (78 mM citric acid, 97 mM sodium citrate, 111 mM D-glucose, pH 7.4) as an anticoagulant. The collected blood was added to a 1.5 mL tube containing 300 μL of HT buffer (129 mM NaCl, 20 mM HEPES, 12 mM NaHCO_3_, 2.9 mM KCl, 1 mM MgCl_2_, 0.34 mM Na_2_HPO_4_, and 0.045 g of glucose per 50 mL of buffer, pH 7.3). Following centrifugation at 250×g for 3 minutes at 25°C, the supernatant (platelet-rich plasma) along with the top one-third of the red cell layer was transferred to a new 1.5 mL tube. This mixture was centrifuged again at 250g for 3 minutes, and the supernatant was collected into another tube. An additional 200 μL of HT buffer was added to the previous tube, which was then centrifuged at 250×g for 3 minutes, and the resulting supernatant was pooled with the previous collection, resulting in approximately 600 μL of platelet-rich plasma. For functional studies, 50 ng/mL of PGI_2_ (Sigma, Cat #P6188, 1 mg/mL in TBS, pH 7.3) was added to the platelet-rich plasma and incubated for 30 minutes at 37°C. The mixture was then centrifuged at 2000×g for 5 minutes to pellet the platelets. All centrifugation steps were performed with slow acceleration and deceleration settings.

### Preparation of platelet-derived extracellular vehicles (PEVs)

Platelets were cultured in RPMI-1640 medium (VivaCell, Cat# C3001-0500) containing 10% exosome-depleted FBS, 100 IU penicillin, and 100 μg/mL streptomycin at 37°C for 2 hours. The culture medium was then collected and centrifuged at 500 g for 10 minutes at 4°C to remove cells. The supernatant was subsequently centrifuged at 2,000×g for 15 minutes at 4°C to remove cell debris. This process was followed by another centrifugation at 10,000×g for 15 minutes at 4°C to eliminate small cell debris. The remaining supernatant was used to isolate PEVs by ultracentrifugation at 30,000 rpm for 90 minutes at 4°C. The obtained PEV pellets were resuspended in an appropriate buffer. PEV-check antibodies TSG101 and CD63 were used to confirm marker expression profiles.

### Transmission Electron Microscopy (TEM)

Isolated PEVs were resuspended in PBS. A 10 μL droplet of the PEV suspension was applied to a formvar-carbon coated copper grid for 1 minutes. The grid was then negatively stained with 2% uranyl acetate for 30 second, air-dried, and visualized using a Hitachi HT7800 transmission electron microscope operating at 120 kV.

### Nanoparticle Tracking Analysis (NTA)

The size distribution and concentration of PEVs were analyzed using a ZetaView (Particle Metrix). PEVs were diluted in sterile PBS to achieve an optimal particle concentration for analysis. Three videos of 60 seconds each were captured for each sample, and the data were processed using ZetaView software (version 9.2).

### In vitro cell culture

For experiments involving KCs, 1×10^^6^ cells seeded in one well of a 12-well plate were co-cultured with either 0.5×10^^6^ isolated mouse platelets or vesicles isolated from 1×10^^6^ isolated mouse platelets for 2 hours. After co-culture, the cells underwent various experiments including Seahorse assay, and RT-PCR. In another study, KCs were treated with platelet supernatant for 2 hours. Platelet supernatant was collected from isolated mouse platelets after a 2-hour culture. In additional experiments, KCs were treated with recombinant ALDOA for 12 hours. In some experiments, cells were treated with either vehicle or 5 μg/mL PEVs for 2 hours.

### Sample preparation for flow cytometry analysis

Hepatic nonparenchymal cells (NPCs) were isolated as described previously(Shan et al., 2021). Briefly, mice were anesthetized and perfused according to the procedure for isolating Kupffer cells. After removing hepatocytes, the remaining cell suspension was centrifuged at 450 × g for 5 minutes to collect NPCs. The pellets were resuspended in 10 mL of 35% Percoll (Cytiva, Cat #17089109) and transferred to a new 15 mL tube. This was followed by centrifugation at 450 x g for 15 minutes to remove dead cells. Isolated liver NPCs were incubated with 1 μL of anti-mouse FcγRII/III (Invitrogen, Cat #14-0161-86) in FACS buffer (PBS + 0.2% FBS) to minimize non-specific antibody binding. The cells were then stained with anti-mouse CD45-APC 780 (Invitrogen, Cat #47-0451-82), F4/80-PE (Invitrogen, Cat #12-4801-82), and CD11b-660 (Invitrogen, Cat #50-0112-82) for 30 minutes on ice. All antibodies were diluted 1:100. Flow cytometry was performed on an LSR Fortessa Cell Analyzer from BD Biosciences, and the data were analyzed using FlowJo software V10.

### Histology and immunofluorescence staining

H&E Staining: Hematoxylin and eosin (H&E) staining was performed on paraffin-embedded liver sections. Briefly, liver tissues were harvested and fixed in 10% formalin for overnight, then dehydrated gradient alcohol. The tissues were subsequently embedded in paraffin and sectioned. H&E staining was performed using Hematoxylin (Biocare Medical, CATHE-M) and Edgar Degas Eosin (Biocare Medical, THE-MM). The necrosis areas were calculated using ImageJ.

Immunofluorescent Staining: Immunofluorescent staining was conducted on frozen liver sections. Fresh liver tissues were harvested and immediately fixed in 4% paraformaldehyde for 1 hour, washed with PBS three times, dehydrated in 30% sucrose at 4°C overnight, and finally embedded in OCT. Frozen sections (5 μm) were prepared and fixed with precooled acetone for 15 minutes at −20°C. The sections were permeabilized with 0.2% Triton X-100 in PBS and blocked with a blocking buffer (5% goat serum + 0.3% BSA in PBS). Following this, the sections were incubated with the primary antibody Purified Rat Anti-Mouse CD41 (BioLegend, Cat #553847, 1:200) overnight at 4°C in a humidified chamber. After washing with TBST, the sections were incubated with Alexa Fluor 488-conjugated Affinipure Goat anti-Rat secondary antibody (Jackson ImmunoResearch, Cat #112-545-003, 1:800) or Alexa Fluor^TM^ 568 goat anti-Rat lgG(H+L) antibody (Invitrogen, Cat #A11077, 1:1000) for 1 hour at room temperature. For Kupffer cells co-staining, sections were further incubated with Alexa Fluor 594 anti-mouse F4/80 (BioLegend, Cat #123140, 1:200) or Alexa Fluor 647 anti-mouse TIM4 (BioLegend, Cat #156804, 1:200) for 1 hour at room temperature. Finally, the sections were stained with DAPI (Beyotime, Cat #C1006) before mounting and imaging on an LSM 900 microscope (Zeiss). All images were collected using consistent parameters.

### DHE staining

Immediately after euthanasia, the liver was excised, rinsed in ice-cold PBS, and cut into small blocks. The tissue blocks were embedded in OCT compound, rapidly frozen on dry ice, and sectioned at 8 µm thickness using a cryostat. The sections were briefly washed twice with PBS, and each tissue section was encircled with a hydrophobic barrier pen. Subsequently, the sections were incubated with 5 µM dihydroethidium (DHE, Servicebio, cat #G1904) at 37 °C for 5 min in a humidified chamber. After two washes with TBST, the sections were mounted with an anti-fade mounting medium and immediately imaged using a fluorescence microscope. The entire procedure from tissue collection to imaging was completed within 1 hour to minimize potential artifacts.

### Sample Preparation for RNA Sequencing and Mass Spectrometry

Mice were treated according to different experimental protocols. KCs were isolated and lysed in 1 mL RNAiso Plus reagent (TaKaRa, Cat #9109). RNA concentration and purity were measured using a NanoDrop 2000 (Thermo Fisher Scientific), and RNA integrity was assessed using the RNA Nano 6000 Assay Kit on the Agilent Bioanalyzer 2100 system (Agilent Technologies). A total of 1 μg RNA per sample was used for RNA sample preparation. Sequencing libraries were generated using the NEB Next Ultra™ RNA Library Prep Kit for Illumina (NEB) following the manufacturer’s recommendations. Index codes were added to attribute sequences to each sample. Clustering of the index-coded samples was performed on a cBot Cluster Generation System using the TruSeq PE Cluster Kit v4-cBot-HS (Illumina) according to the manufacturer’s instructions. After cluster generation, the library preparations were sequenced on an Illumina NovaSeq 6000 platform, generating paired-end reads (the sequencing data for Fig1A-B were performed in BGI, and the sequencing data for Fig1C-D were performed in BMKCloud).

### Mass Spectrometry Analysis of EV Cargo

To identify PEV-specific protein cargo, PEVs were isolated as described. For comparison, hepatocyte-derived extracellular vesicles (HEVs) were isolated from the culture supernatant of AML12, a mouse hepatocytes cell line, using an identical ultracentrifugation protocol. Protein extracts from PEVs and HEVs were trypsin-digested and analyzed by liquid chromatography-tandem mass spectrometry (LC-MS/MS) on a Orbitrap Exploris 480 MS (Thermo Fisher Scientific). Proteins identified in HEVs, as well as those annotated to the Gene Ontology (GO) terms “cytoskeleton” (GO:0005856) and “blood coagulation” (GO:0007596), were subtracted from the PEV proteome. Candidate proteins were defined as those present in all three biological replicates with more than two unique peptides.

### RNA sequencing analysis

Raw data (raw reads) in fastq format were processed using in-house Perl scripts. Clean data (clean reads) were obtained by removing reads containing adapters, reads containing poly-N, and low-quality reads from the raw data. Quality metrics such as Q20, Q30, GC-content, and sequence duplication levels were calculated for the clean data. All downstream analyses were based on high-quality clean data. Reads were mapped against the reference genome and transcript annotation (GRCm38) using Hisat2 software. Further analysis was performed using RStudio (version 4.3). Differentially expressed genes were determined using the limma package, and enrichment analysis was conducted using clusterProfiler.

### Western blot

Cells were lysed in RIPA lysis buffer (10mM Tris-HCl pH 8.0, 1mM SDTA, 0.5mM EGTA, 1% Triton X-100, 0.1%Ssodium deoxycholate, 0.1% SDS, 140mM NaCl, 100mM PMSF and 100 ×Cocktail were added freshly when using) on ice for 30 minutes. Protein extracts were obtained from the supernatants following centrifugation and subjected to SDS-PAGE. The membranes were blocked with 5% non-fat dry milk for 1 hour at room temperature. After blocking, membranes were incubated with primary antibodies followed by species-appropriate horseradish peroxidase-conjugated secondary antibodies. Membranes were washed three times with TBS-T and detected using either ECL Select Western Blotting Detection Reagent (Cytiva, Cat #RPN2235) or SuperSignal West Pico PLUS Chemiluminescent Substrate (Thermo Scientific, Cat #34580). For mouse serum, which will be diluted with PBS for 20 times, and then add SDS-loading buffer, 98℃ for 10 min.

Primary antibodies used to detect specific proteins: ALDOA Polyclonal antibody (Proteintech, Cat #11217-1-AP, 1:5000), TSG101 antibody(C-2) (Santa Cruz, Cat #sc-7964), Beta Actin Monoclonal antibody (Proteintech, Cat #66009-1-Ig, 1:10,000), Albumin Antibody (Cell Signaling Technology, Cat #4929, 1:1000), Alpha-Tubulin monoclonal antibody (Proteintech, 66031-1-Ig, 1:2000). Secondary antibodies include goat anti-Rabbit IgG (Jackson ImmunoResearch, Cat #111-035-003, 1:10,000), goat anti-Mouse IgG (Jackson ImmunoResearch, Cat #115-035-003, 1:10,000).

### GSH measurement

The GSH detection was performed according to the protocol of the official kit (Reduced Glutathione (GSH) Content Assay Kit, Solarbio, cat #BC1175). Briefly, liver tissues were collected, snap-frozen in liquid nitrogen, and ground. Then, 30 mg of tissue was weighed and mixed with 300 μL of Reagent 1. After ultrasonic disruption, the samples were centrifuged at 12,000 rpm for 10 min at 4°C. The supernatant was transferred to a new 1.5 mL EP tube. In a 96-well plate, standards (0, 31.25, 62.5, 125, 250, 500, 1000 μg/mL), 20 μL sample or standards, 140 μL Reagent 2, and 40 μL Reagent 3 were added. The absorbance was measured at OD412 nm using a microplate reader. Meanwhile, the protein concentration of the samples was determined using a BCA kit. Finally, the GSH concentration was normalized to the protein concentration.

### Seahorse Analysis

Real-time changes in extracellular acidification rate (ECAR) of KCs were measured using the XFe24 Seahorse extracellular flux analyzer (Agilent) with the Seahorse XF Glycolysis Stress Test Kit (Agilent, Cat #103020-100). KCs were plated onto XFe24 cell culture plates containing 0.3×10^6 cells a well and subjected to specific experimental treatments. The cells were rinsed with Seahorse XF basal medium supplemented with 2 mM glutamine and maintained in 500 µL of the same basal medium. The analysis was performed using the following Seahorse XF running buffers: Buffer A: 100 mM glucose in basal medium;Buffer B: 10 μM oligomycin in basal medium;Buffer C: 500 mM 2-deoxy-D-glucose (2-DG) in basal medium. At the end of the assay, the cells were lysed with RIPA buffer, and protein quantification was performed using the Pierce™ Protein Assay Kit (Thermo Scientific, Cat #23227) based on the bicinchoninic acid (BCA) method. Results were analyzed using Seahorse Wave Desktop Software (Agilent) and were normalized to protein content.

### Prokaryotic purification of recombinant ALDOA

For the prokaryotic purification of recombinant Aldoa, the full-length cDNAs encoding Aldoa were cloned into the pET-28a vector and transformed into E. coli strain BL21 (DE3). The transformed cells were induced with 1 mM isopropyl-β-D-thiogalactopyranoside at an OD600 of 1.0 and incubated for an additional 4 hours at 37°C. Following incubation, the cells were collected and resuspended in ice-cold His binding buffer (20 mM Tris-HCl, pH 8.0, 500 mM NaCl, 1% Triton X-100, 1% glycerol, 5 mM imidazole). The cell suspension was sonicated and then centrifuged at 15,000 g for 1 hour at 4°C. The supernatant was filtered through a 40 μm filter membrane and subsequently purified using Ni-NTA Agarose. The Ni-NTA Agarose was washed sequentially with ice-cold His wash buffers: first with buffer 1 (20 mM Tris-HCl, pH 8.0, 500 mM NaCl, 1% glycerol, 20 mM imidazole), followed by buffer 2 (20 mM Tris-HCl, pH 8.0, 500 mM NaCl, 1% glycerol, 40 mM imidazole). Recombinant Aldoa were eluted from the resin using ice-cold His elution buffer (20 mM Tris-HCl, pH 8.0, 500 mM NaCl, 1% glycerol, 500 mM imidazole) and its concentration was determined using a BCA assay.

### Real-time RT-PCR assay

Total RNA was extracted from cells using RNAiso Plus reagent, followed by chloroform extraction and isopropanol precipitation to isolate RNA. The RNA pellets were washed with 75% ethanol, air-dried, and resuspended in DEPC-treated water. cDNA synthesis was carried out using the PrimeScript II 1st Strand cDNA Synthesis Kit (TaKaRa, Cat #6210A), adhering to the manufacturer’s protocols. Quantitative RT-PCR (qPCR) was performed in duplicate using the PowerUp SYBR Green Master Mix (Applied Biosystems, Cat #A25742) on a StepOne system. The PCR conditions included an initial denaturation at 50°C for 2 minutes, followed by 95°C for 2 minutes, and then 40 cycles of 95°C for 15 seconds and 60°C for 1 minute. A final cycle involved 95°C for 15 seconds, 60°C for 1 minute, and 95°C for 15 seconds. Data were analyzed using the comparative Ct method with 18S rRNA as the internal control. Primer sequences are detailed in Supplementary File 3.

### Analysis of Human Proteomics Data

For the analysis of human platelet alpha granules (Fig 6F), the raw mass spectrometry data from Maynard et al. were obtained and re-analyzed using Graphpad Prism software (v.8.0.2) to extract the spectral counts for ALDOA. For the analysis of EV proteomes from polycythemia vera and Alzheimer’s disease patients (Fig 6G), processed data tables from Fel et al (PXD013234) and Nielsen et al (PXD024216) were directly obtained from the ProteomeXchange database as provided in the respective publications, and the normalized intensity values for ALDOA were extracted.

### Statistical analysis

Statistical analyses were conducted using GraphPad Prism software (v.8.0.2). Data are expressed as mean values ± SEM. To assess significance between two groups, an unpaired two-tailed Student’s t-test was employed. For comparisons among multiple groups, one-way ANOVA was used. Statistical significance was defined as *p* < 0.05. Detailed statistical information is provided in the figure legends and associated data. Images shown without biological replicates represent results from at least three independent experiments.

## Data and code availability

RNA-seq data for this study have been deposited in Gene Expression Omnibus at National Center for Biotechnology Information accession number GSE274627.

## Author Contributions

R.X.Y. conducted the experiments, analysed the data, and wrote the manuscript. J.H.L. provided the patient samples. T.W., Y.H.L., C.S., and L.Y. were responsible for patient sample collection. K.F. performed the transmission electron microscopy analysis of PEVs. K.Q.W. prepared the mass spectrometry samples of HEVs. Z.S. conceived and designed the study, supervised the project, and contributed to the writing and revision of the manuscript.

## Acknowledgements

We thank Dr. Bin Qi (Yunnan University) and Dr. Wenxiang Fu (Yunnan University) for suggestions and discussions. We thank Dr. Hao Yin (Wuhan University) for providing us with AML12 cells.

## Additional information

### Data availability

All data generated or analysed during this study are included in the manuscript and supporting files; source data files have been provided.

## Supplementary File

**Supplementary File 1** PEVs MS

**Supplementary File 2** Clinical characteristics of patients with ALI

**Supplementary File 3** Real-Time PCR Primers used

