## Supplemental figure for "Platelets promote acute liver injury via extracellular vesicles-mediated Aldolase A"

### Slide 1
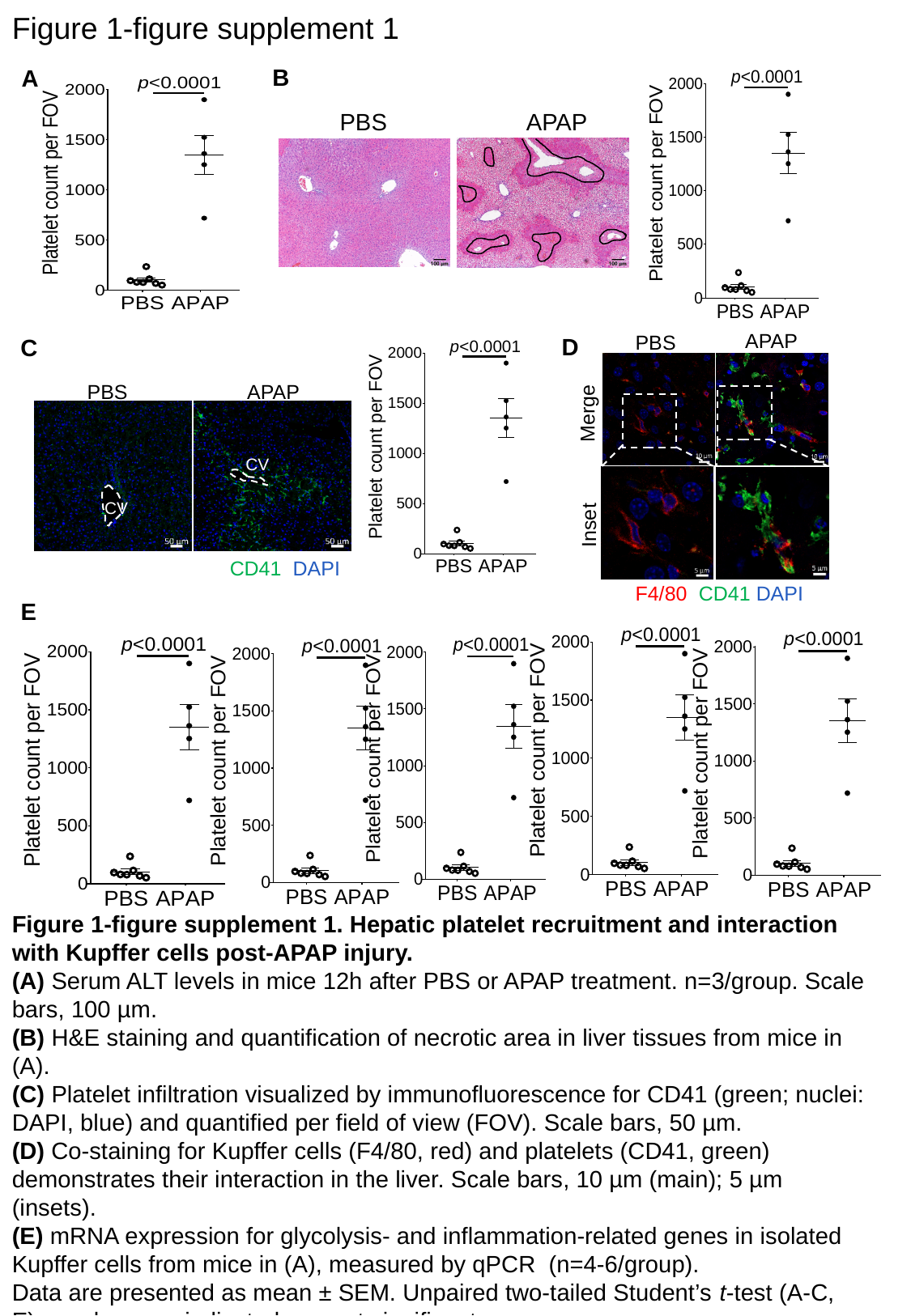

Figure 1-figure supplement 1
B
A
APAP
PBS
APAP
PBS
Merge
Inset
F4/80 CD41 DAPI
D
C
PBS
APAP
CD41 DAPI
CV
CV
E
Figure 1-figure supplement 1. Hepatic platelet recruitment and interaction with Kupffer cells post-APAP injury.
(A) Serum ALT levels in mice 12h after PBS or APAP treatment. n=3/group. Scale bars, 100 µm.
(B) H&E staining and quantification of necrotic area in liver tissues from mice in (A).
(C) Platelet infiltration visualized by immunofluorescence for CD41 (green; nuclei: DAPI, blue) and quantified per field of view (FOV). Scale bars, 50 µm.
(D) Co-staining for Kupffer cells (F4/80, red) and platelets (CD41, green) demonstrates their interaction in the liver. Scale bars, 10 µm (main); 5 µm (insets).
(E) mRNA expression for glycolysis- and inflammation-related genes in isolated Kupffer cells from mice in (A), measured by qPCR (n=4-6/group).
Data are presented as mean ± SEM. Unpaired two-tailed Student’s t-test (A-C, E). p values are indicated; ns, not significant.

### Slide 2
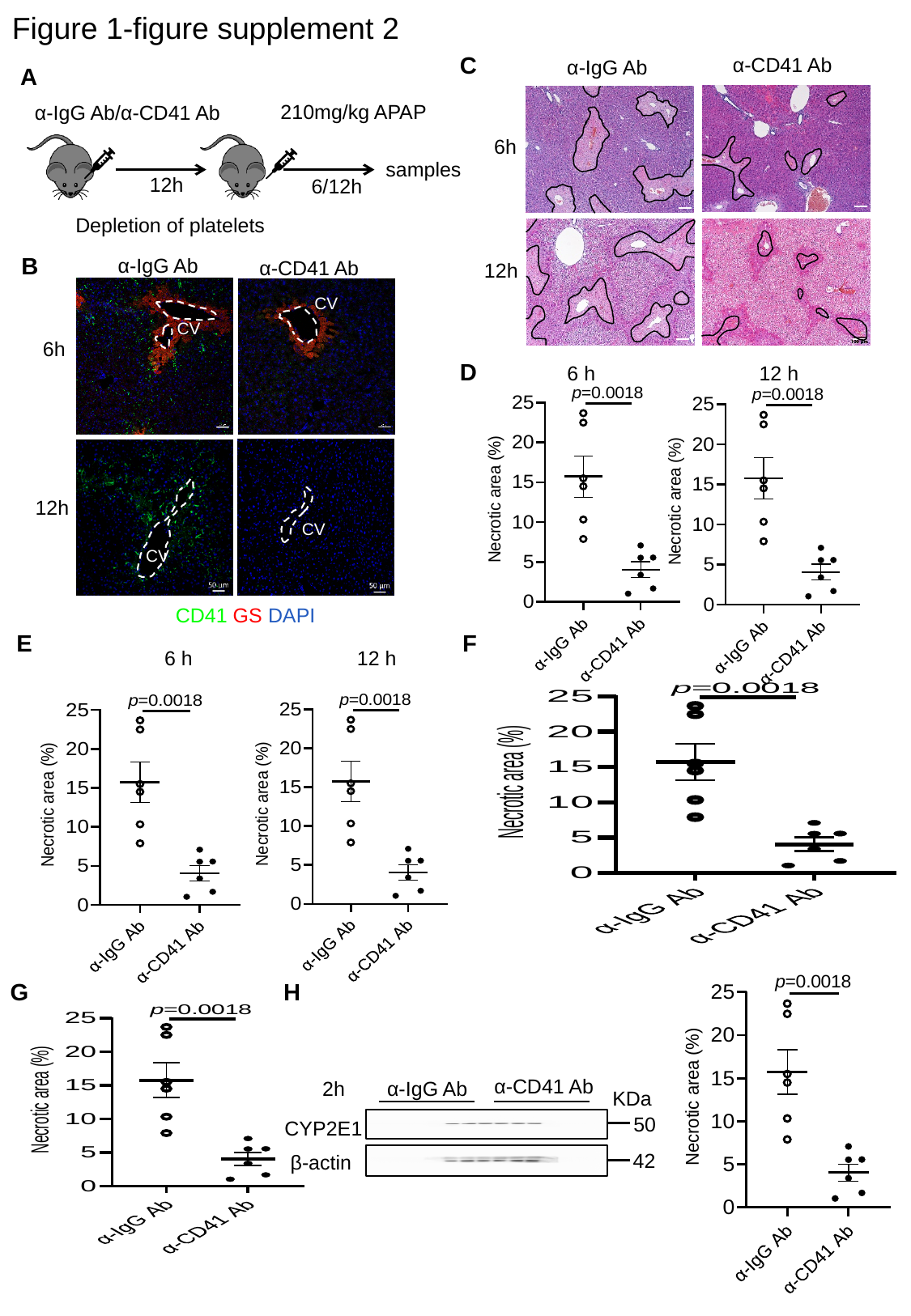

Figure 1-figure supplement 2
C
α-CD41 Ab
α-IgG Ab
A
210mg/kg APAP
samples
6/12h
α-IgG Ab/α-CD41 Ab
12h
Depletion of platelets
6h
B
α-IgG Ab
α-CD41 Ab
CD41 GS DAPI
CV
CV
6h
12h
CV
CV
12h
D
6 h
12 h
E
F
6 h
12 h
G
H
α-CD41 Ab
α-IgG Ab
2h
50
CYP2E1
42
β-actin
KDa

### Slide 3
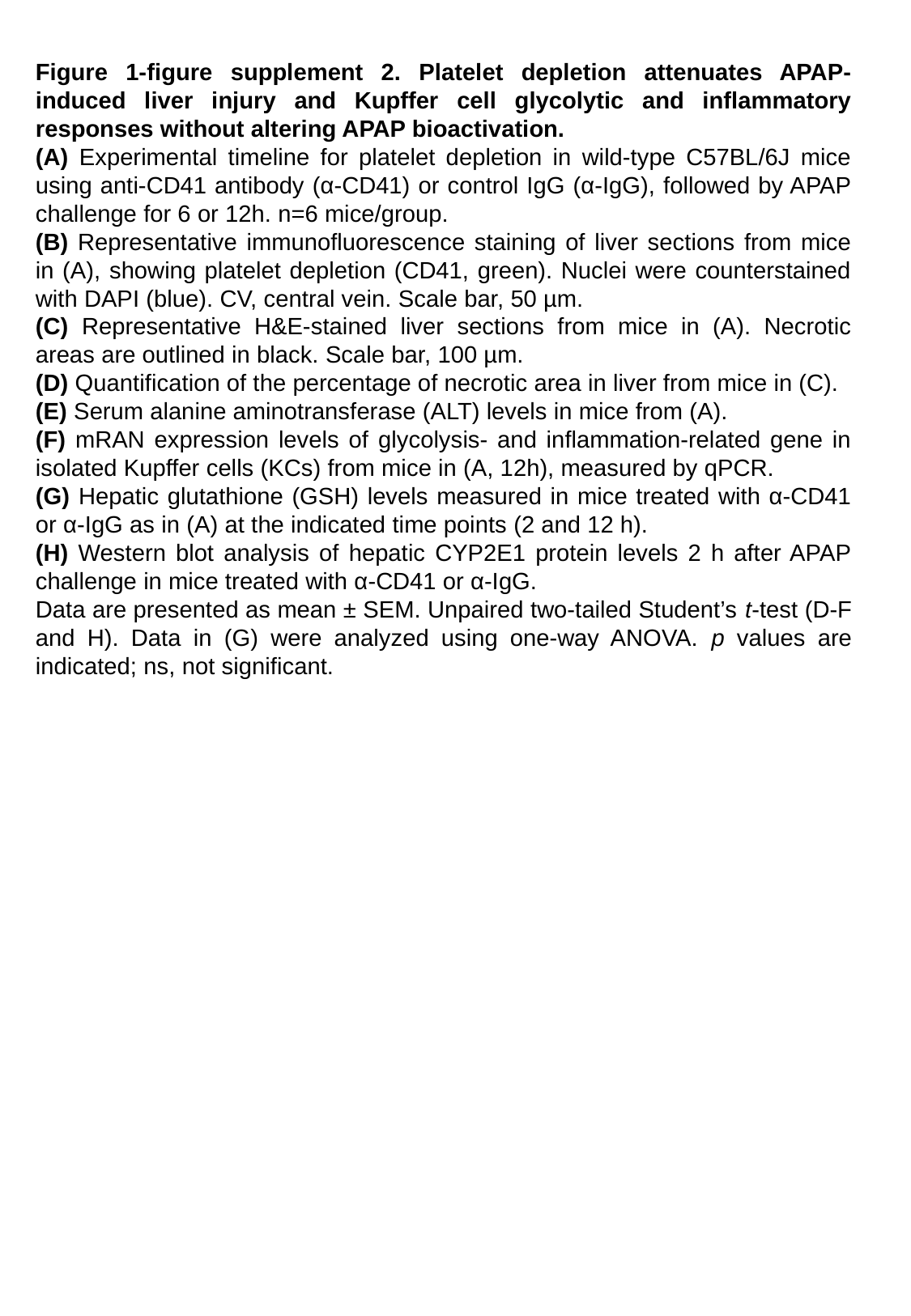

Figure 1-figure supplement 2. Platelet depletion attenuates APAP-induced liver injury and Kupffer cell glycolytic and inflammatory responses without altering APAP bioactivation.
(A) Experimental timeline for platelet depletion in wild-type C57BL/6J mice using anti-CD41 antibody (α-CD41) or control IgG (α-IgG), followed by APAP challenge for 6 or 12h. n=6 mice/group.
(B) Representative immunofluorescence staining of liver sections from mice in (A), showing platelet depletion (CD41, green). Nuclei were counterstained with DAPI (blue). CV, central vein. Scale bar, 50 µm.
(C) Representative H&E-stained liver sections from mice in (A). Necrotic areas are outlined in black. Scale bar, 100 µm.
(D) Quantification of the percentage of necrotic area in liver from mice in (C).
(E) Serum alanine aminotransferase (ALT) levels in mice from (A).
(F) mRAN expression levels of glycolysis- and inflammation-related gene in isolated Kupffer cells (KCs) from mice in (A, 12h), measured by qPCR.
(G) Hepatic glutathione (GSH) levels measured in mice treated with α-CD41 or α-IgG as in (A) at the indicated time points (2 and 12 h).
(H) Western blot analysis of hepatic CYP2E1 protein levels 2 h after APAP challenge in mice treated with α-CD41 or α-IgG.
Data are presented as mean ± SEM. Unpaired two-tailed Student’s t-test (D-F and H). Data in (G) were analyzed using one-way ANOVA. p values are indicated; ns, not significant.

### Slide 4
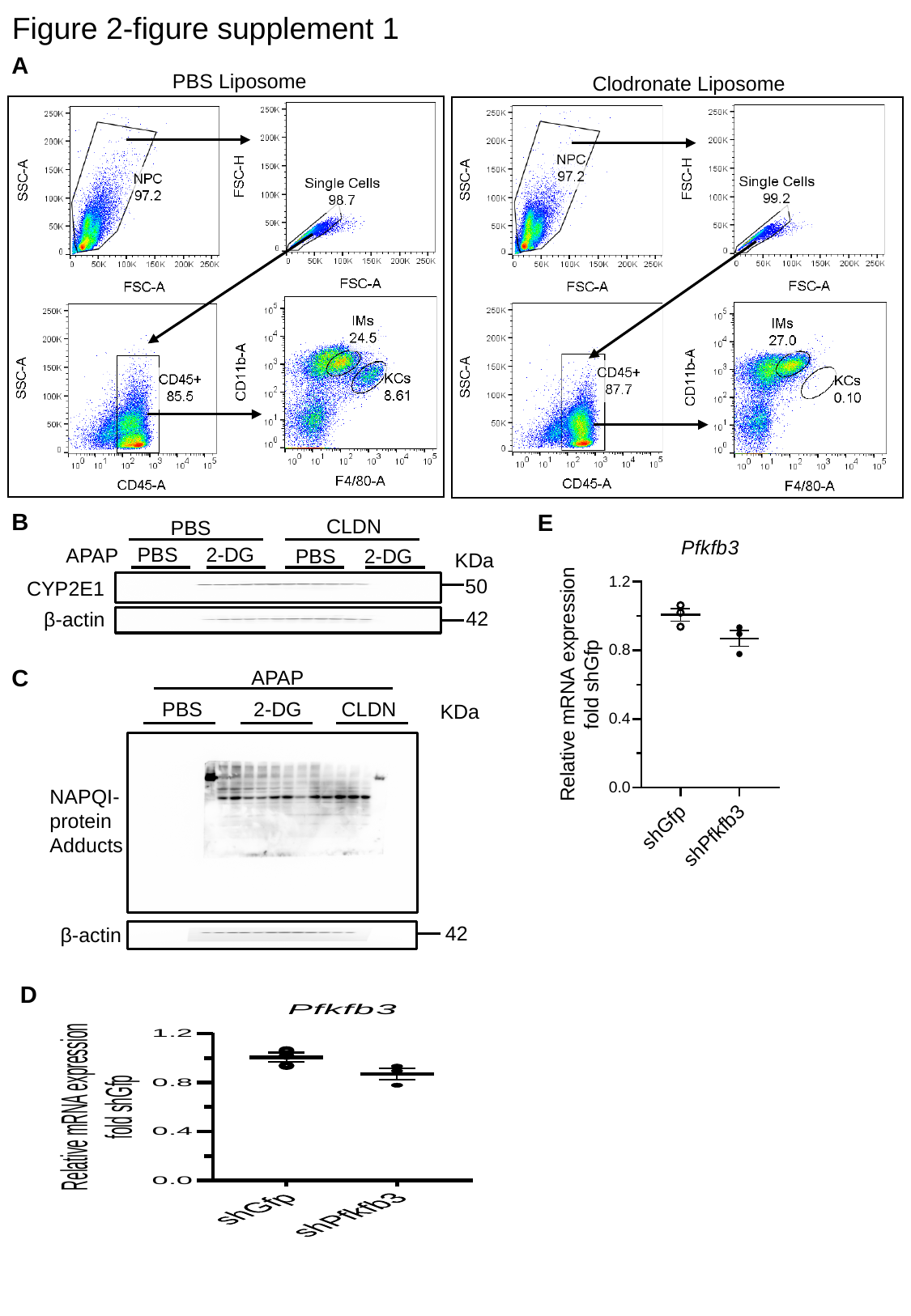

Figure 2-figure supplement 1
A
PBS Liposome
Clodronate Liposome
B
E
CLDN
PBS
PBS 2-DG
50
CYP2E1
42
β-actin
APAP
PBS 2-DG
KDa
C
APAP
PBS 2-DG CLDN
KDa
NAPQI-protein Adducts
42
β-actin
D

### Slide 5
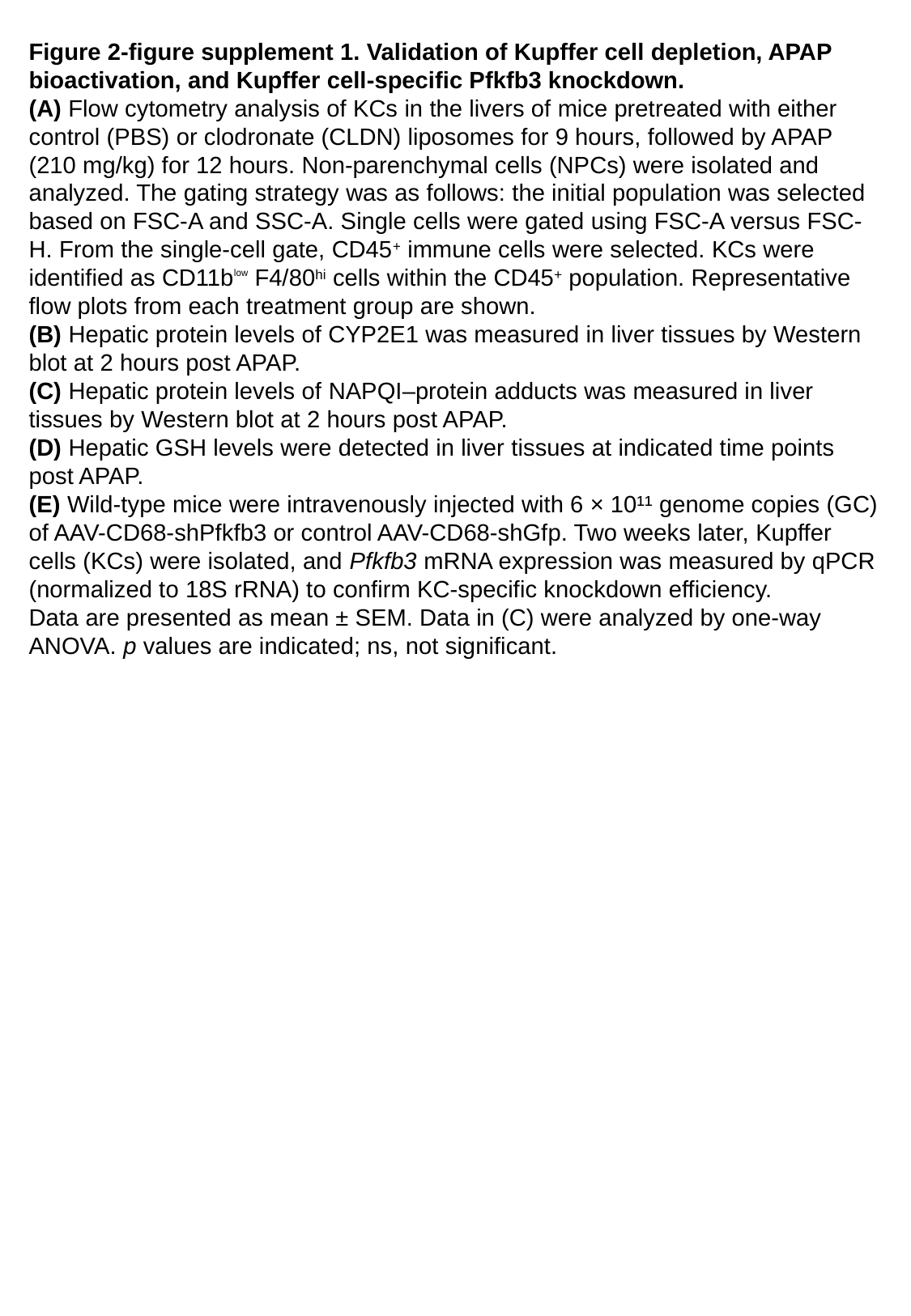

Figure 2-figure supplement 1. Validation of Kupffer cell depletion, APAP bioactivation, and Kupffer cell-specific Pfkfb3 knockdown.
(A) Flow cytometry analysis of KCs in the livers of mice pretreated with either control (PBS) or clodronate (CLDN) liposomes for 9 hours, followed by APAP (210 mg/kg) for 12 hours. Non-parenchymal cells (NPCs) were isolated and analyzed. The gating strategy was as follows: the initial population was selected based on FSC-A and SSC-A. Single cells were gated using FSC-A versus FSC-H. From the single-cell gate, CD45^+^ immune cells were selected. KCs were identified as CD11b^ˡᵒʷ^ F4/80^hi^ cells within the CD45^+^ population. Representative flow plots from each treatment group are shown.
(B) Hepatic protein levels of CYP2E1 was measured in liver tissues by Western blot at 2 hours post APAP.
(C) Hepatic protein levels of NAPQI–protein adducts was measured in liver tissues by Western blot at 2 hours post APAP.
(D) Hepatic GSH levels were detected in liver tissues at indicated time points post APAP.
(E) Wild-type mice were intravenously injected with 6 × 10¹¹ genome copies (GC) of AAV-CD68-shPfkfb3 or control AAV-CD68-shGfp. Two weeks later, Kupffer cells (KCs) were isolated, and Pfkfb3 mRNA expression was measured by qPCR (normalized to 18S rRNA) to confirm KC-specific knockdown efficiency.
Data are presented as mean ± SEM. Data in (C) were analyzed by one-way ANOVA. p values are indicated; ns, not significant.

### Slide 6
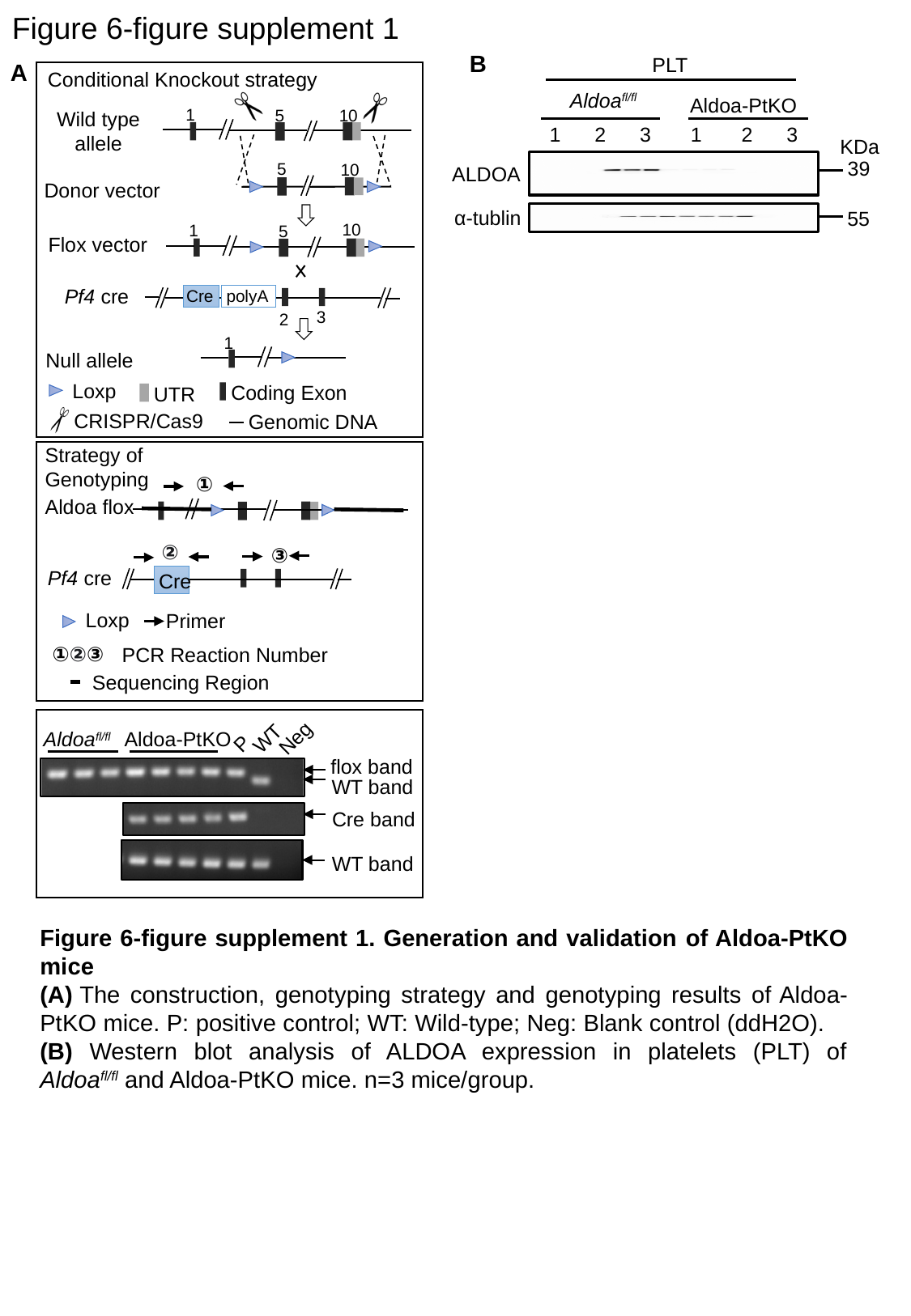

Figure 6-figure supplement 1
B
PLT
Aldoa^fl/fl^
Aldoa-PtKO
39
ALDOA
α-tublin
55
1 2 3 1 2 3
KDa
A
Conditional Knockout strategy
1
10
5
Wild type allele
5
10
Donor vector
10
1
5
Flox vector
Pf4 cre
Cre
1
Null allele
Loxp
Coding Exon
UTR
CRISPR/Cas9
Genomic DNA
polyA
3
2
Strategy of Genotyping
①
Aldoa flox
②
③
Cre
Loxp
Primer
①②③
PCR Reaction Number
Sequencing Region
Pf4 cre
P
Neg
WT
Aldoa^fl/fl^
Aldoa-PtKO
flox band
WT band
Cre band
WT band
Figure 6-figure supplement 1. Generation and validation of Aldoa-PtKO mice
(A) The construction, genotyping strategy and genotyping results of Aldoa-PtKO mice. P: positive control; WT: Wild-type; Neg: Blank control (ddH2O).
(B) Western blot analysis of ALDOA expression in platelets (PLT) of Aldoa^fl/fl^ and Aldoa-PtKO mice. n=3 mice/group.

### Slide 7
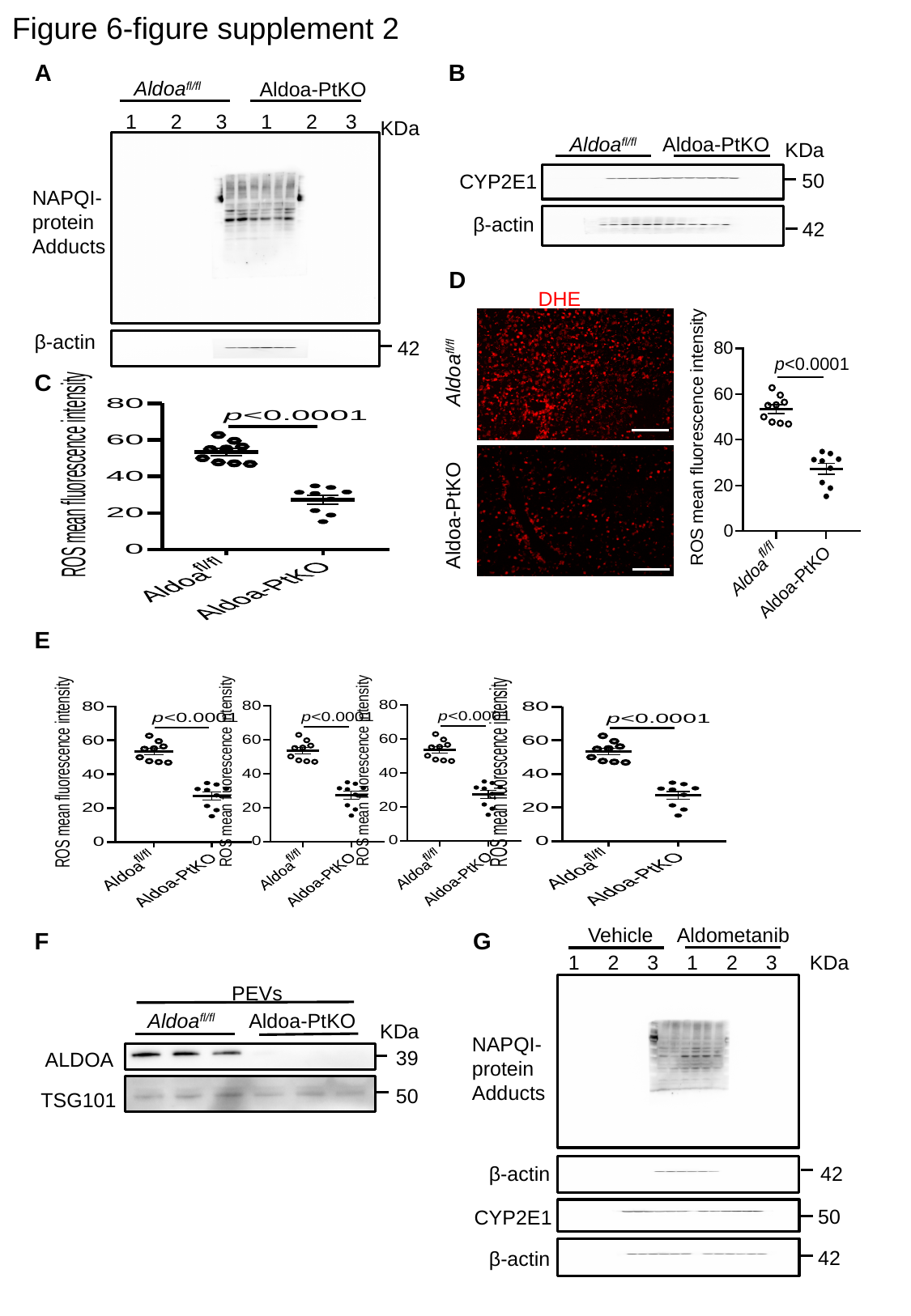

Figure 6-figure supplement 2
A
B
Aldoa^fl/fl^
Aldoa-PtKO
1 2 3 1 2 3
42
NAPQI-protein Adducts
β-actin
KDa
Aldoa^fl/fl^ Aldoa-PtKO
KDa
50
CYP2E1
β-actin
42
D
DHE
Aldoa-PtKO Aldoa^fl/fl^
C
E
Vehicle
Aldometanib
1 2 3 1 2 3
NAPQI-protein Adducts
42
β-actin
F
G
KDa
PEVs
Aldoa^fl/fl^ Aldoa-PtKO
KDa
39
ALDOA
50
TSG101
50
CYP2E1
42
β-actin

### Slide 8
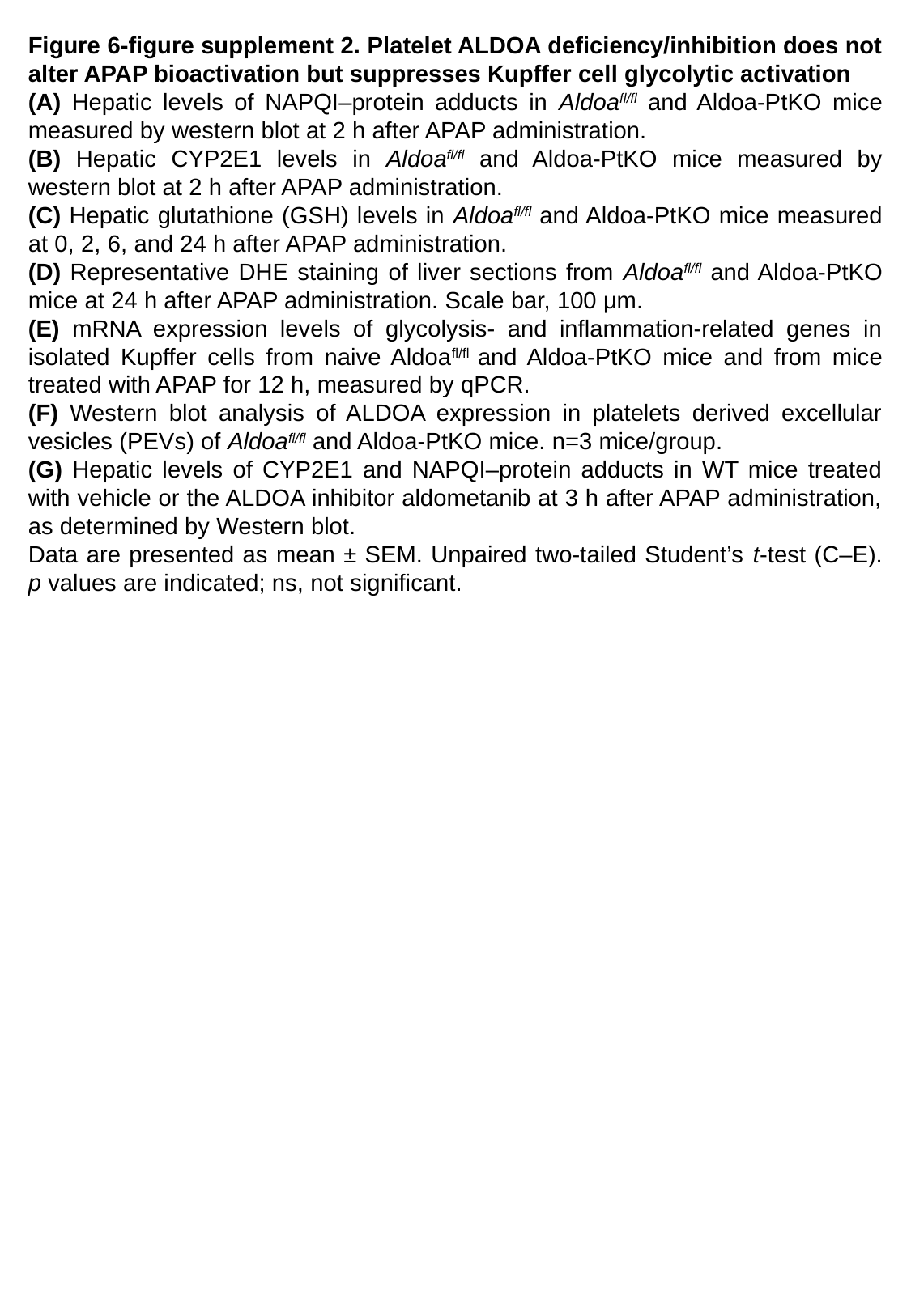

Figure 6-figure supplement 2. Platelet ALDOA deficiency/inhibition does not alter APAP bioactivation but suppresses Kupffer cell glycolytic activation
(A) Hepatic levels of NAPQI–protein adducts in Aldoa^fl/fl^ and Aldoa-PtKO mice measured by western blot at 2 h after APAP administration.
(B) Hepatic CYP2E1 levels in Aldoa^fl/fl^ and Aldoa-PtKO mice measured by western blot at 2 h after APAP administration.
(C) Hepatic glutathione (GSH) levels in Aldoa^fl/fl^ and Aldoa-PtKO mice measured at 0, 2, 6, and 24 h after APAP administration.
(D) Representative DHE staining of liver sections from Aldoa^fl/fl^ and Aldoa-PtKO mice at 24 h after APAP administration. Scale bar, 100 μm.
(E) mRNA expression levels of glycolysis- and inflammation-related genes in isolated Kupffer cells from naive Aldoa^fl/fl^ and Aldoa-PtKO mice and from mice treated with APAP for 12 h, measured by qPCR.
(F) Western blot analysis of ALDOA expression in platelets derived excellular vesicles (PEVs) of Aldoa^fl/fl^ and Aldoa-PtKO mice. n=3 mice/group.
(G) Hepatic levels of CYP2E1 and NAPQI–protein adducts in WT mice treated with vehicle or the ALDOA inhibitor aldometanib at 3 h after APAP administration, as determined by Western blot.
Data are presented as mean ± SEM. Unpaired two-tailed Student’s t-test (C–E). p values are indicated; ns, not significant.

### Slide 9
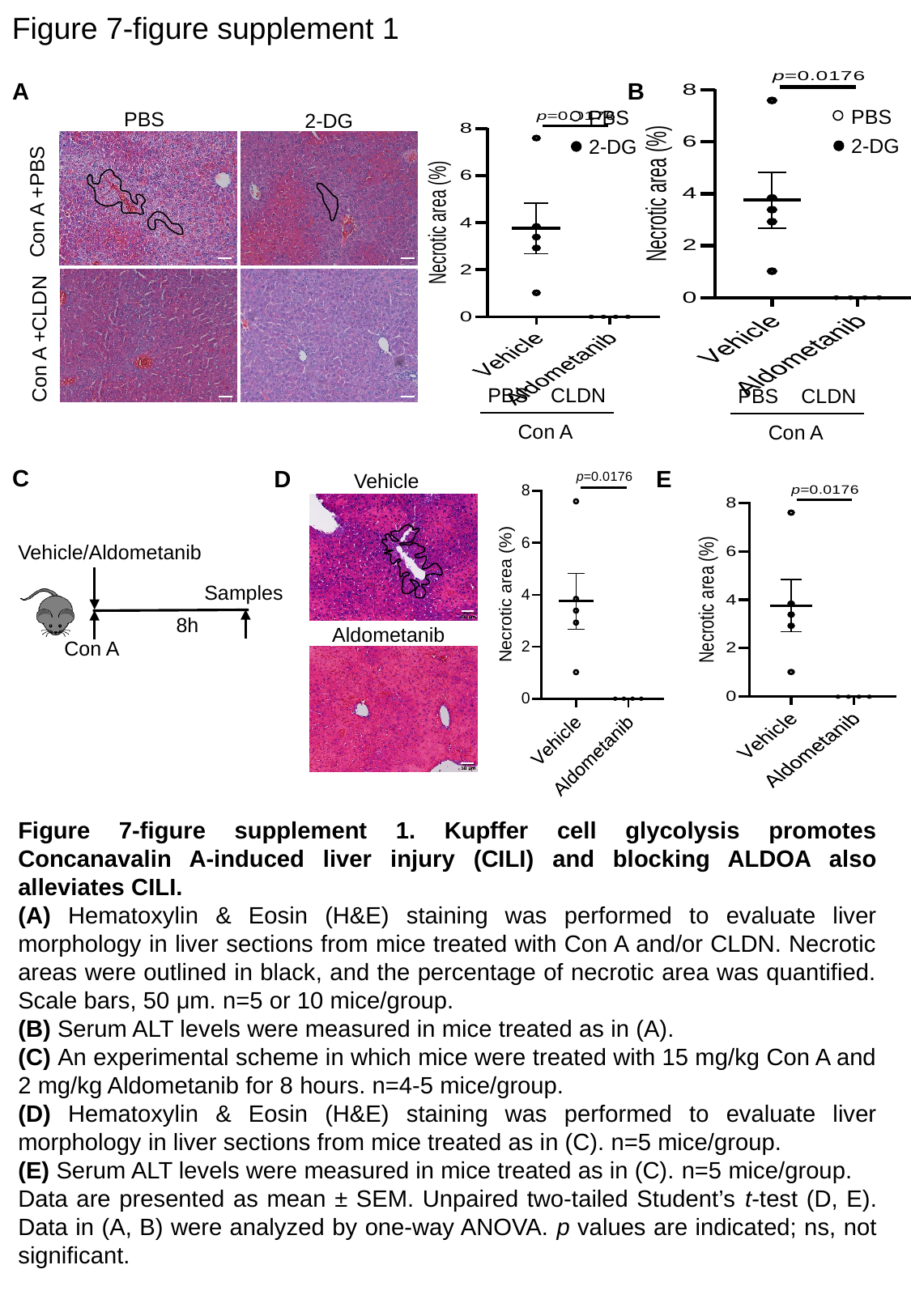

Figure 7-figure supplement 1
B
A
PBS
2-DG
PBS
2-DG
PBS CLDN
Con A
PBS
2-DG
Con A +PBS
Con A +CLDN
PBS CLDN
Con A
Vehicle
Aldometanib
C
E
D
Vehicle/Aldometanib
Samples
Con A
8h
Figure 7-figure supplement 1. Kupffer cell glycolysis promotes Concanavalin A-induced liver injury (CILI) and blocking ALDOA also alleviates CILI.
(A) Hematoxylin & Eosin (H&E) staining was performed to evaluate liver morphology in liver sections from mice treated with Con A and/or CLDN. Necrotic areas were outlined in black, and the percentage of necrotic area was quantified. Scale bars, 50 μm. n=5 or 10 mice/group.
(B) Serum ALT levels were measured in mice treated as in (A).
(C) An experimental scheme in which mice were treated with 15 mg/kg Con A and 2 mg/kg Aldometanib for 8 hours. n=4-5 mice/group.
(D) Hematoxylin & Eosin (H&E) staining was performed to evaluate liver morphology in liver sections from mice treated as in (C). n=5 mice/group.
(E) Serum ALT levels were measured in mice treated as in (C). n=5 mice/group.
Data are presented as mean ± SEM. Unpaired two-tailed Student’s t-test (D, E). Data in (A, B) were analyzed by one-way ANOVA. p values are indicated; ns, not significant.
