## Supplemental File 2 for "Platelets promote acute liver injury via extracellular vesicles-mediated Aldolase A"

| **Table S3 Clinical characteristics of the acute liver injury patients included in this study** | | | | | |
| --- | --- | --- | --- | --- | --- |
| **NO.** | **Gender (male/female)** | **Age (years)** | **Etiology** | **ALT (U/L)** | **TBiL (μmol/L)** |
| 1 | male | 45 | Acute Toxic Liver Injury | 1629 | 26.9 |
| 2 | female | 19 | Acute drug-induced liver injury | 395 | 14.6 |
| 3 | male | 27 | Acute Pesticide Poisoning | 395 | 14.6 |
| 4 | male | 67 | Amatoxin Poisoning | 116 | 30.2 |
| 5 | male | 52 | Amatoxin Poisoning | 536 | 14.9 |
| 6 | female | 41 | Amatoxin Poisoning | 206 | 25.4 |
| 7 | male | 29 | Amatoxin Poisoning | 49.7 | 11.6 |
| 8 | male | 34 | Amatoxin Poisoning | 293 | 23.6 |
| 9 | female | 55 | Amatoxin Poisoning | 3183 | 57.3 |
| 10 | female | 30 | Acute Drug Intoxication (Quetiapine, Olanzapine, and Estazolam) | 127 | 26.1 |
| 11 | male | 30 | Acute Pesticide Poisoning | 261 | 11 |
| 12 | female | 28 | Amatoxin Poisoning | 491 | 19.5 |
| 13 | male | 55 | Amatoxin Poisoning | 572 | 20.3 |
| 14 | male | 65 | Amatoxin Poisoning | 563 | 22.4 |
| 15 | female | 58 | Amatoxin Poisoning | 1154 | 14.7 |
| 16 | male | 75 | Acute drug-induced liver failure | 1240 | 43.1 |

ALT: Alanine Transaminase; TBiL: Total Bilirubin
